# Engineered human iPS cell models reveal altered podocytogenesis and glomerular capillary wall in CHD-associated *SMAD2* mutations

**DOI:** 10.1101/2024.08.02.606108

**Authors:** Rohan Bhattacharya, Tarsha Ward, Titilola D. Kalejaiye, Alekshyander Mishra, Sophia Leeman, Hamidreza Arzaghi, Jonathan G. Seidman, Christine E. Seidman, Samira Musah

**Affiliations:** Department of Biomedical Engineering, Pratt School of Engineering, Duke University, Durham, North Carolina 27705, USA; Center for Biomolecular and Tissue Engineering, Duke University, Durham, North Carolina 27705, USA; Department of Genetics, Harvard Medical School, Boston, Massachusetts 02115, USA; Cardiovascular Division, Department of Medicine, Brigham and Women’s Hospital, Boston, Massachusetts 02115, USA; Howard Hughes Medical Institute (HHMI), Harvard University, Boston, Massachusetts 02115, USA; Division of Nephrology, Department of Medicine, Duke University School of Medicine, Durham, North Carolina 27705, USA; Developmental and Stem Cell Biology Program, Duke University, Durham, North Carolina 27705, USA; Department of Cell Biology, Duke University, Durham, North Carolina, USA; Current affiliation: Boston Children’s Hospital, Harvard Medical School, Boston, Massachusetts 02115, USA

## Abstract

Early developmental programming involves extensive cell lineage diversification through shared molecular signaling networks. Clinical observations of congenital heart disease (CHD) patients carrying *SMAD2* genetic variants revealed correlations with multi-organ impairments at the developmental and functional levels. For example, many CHD patients present with glomerulosclerosis, periglomerular fibrosis, and albuminuria. Still, it remains largely unknown whether *SMAD2* variants associated with CHD can directly alter kidney cell fate, tissue patterning, and organ-level function. To address this question, we engineered human iPS cells (iPSCs) and organ-on-a-chip systems to uncover the role of pathogenic *SMAD2* variants in kidney podocytogenesis. Our results show that abrogation of *SMAD2* causes altered patterning of the mesoderm and intermediate mesoderm (IM) cell lineages, which give rise to nearly all kidney cell types. Upon further differentiation of IM cells, the mutant podocytes failed to develop arborizations and interdigitations. A reconstituted glomerulus-on-a-chip platform exhibited significant proteinuria as clinically observed in glomerulopathies. This study implicates CHD-associated *SMAD2* mutations in kidney tissue malformation and provides opportunities for therapeutic discovery in the future.

---

The development and patterning of human heart and kidneys is highly orchestrated and relies on shared molecular signaling pathways^1,2^. Consequently, mutations in genetic factors involved in organ patterning can lead to multiple congenital defects in the heart and extracardiac targets^3–6^. In the United States, Congenital Heart Disease (CHD) is a common cause of childhood mortality and morbidity that affect 1 in 1000 births^7,8^, and extracardiac anomalies occur in 25% of those patients^9–13^. Although advances in surgical procedures are improving CHD outcomes, widespread extracardiac congenital anomalies can severely compromise life expectancy^14–18^. Investigating how pathogenic mutations associated with CHD cause extracardiac anomalies could advance current understanding of developmentally regulated mechanisms, tissue patterning, and inform therapeutic strategies.

Several developmental abnormalities have been observed in CHD patients who harbor LoF variants in CHD-causing genes^19^. CHD patients often exhibit renal abnormalities, but causal relationships between CHD and extracardiac complications is less understood^9–11,14–16^. Congenital anomalies of the kidneys can affect 1 in 500 newborns and constitute 20–30% of all anomalies identified in the prenatal period^20–22^. However, it remains unclear whether genetic variants linked to CHD can directly alter renal developmental programming and tissue patterning.

Recent reports show LoF mutations in the *SMAD2* gene have been identified in CHD patients having renal abnormalities^23^. Patients with CHD often have hypertension^24^ and hypertensive patients can have 46% less glomeruli^25^. Patients with pathogenic *SMAD2* mutation can have multiple bilateral kidney cysts^23^ and renal cysts account for a 2.3-fold increased prevalence of albuminuria and renal insufficiency^26–28^. The multi-organ defect commonly observed in patients highlights the disease severity associated with *SMAD2* mutations. Additionally, *SMAD2* mutations identified in CHD patients of Pediatric congenital consortium (PCGC) (ClinicalTrials. gov identifier NCT01196182)^12,29,30^ have ureteral defects (stenosis), genitourinary abnormalities, and hydronephrosis, which are known to impact the kidney’s blood filtration function^26–28^. The association of glomerular filtration dysfunction in patients with CHD dates back to several decades, when Boichis^31^ and Guignard^32^ observed leaky filtration barrier in pediatric CHD patients. While many patients exhibit renal dysfunction even before they undergo cardiac repair surgery^33^, almost all patients develop Acute Kidney Injury (AKI) and, subsequently, Chronic Kidney Disease (CKD) after surgery^11,33–39^. Chromosomal rearrangements, including those of 18q21, a region where *SMAD2* maps, are associated with horseshoe kidney^40^, hydronephrosis and fibrosis^41^, cystic kidneys, and ureteral agenesis^42^. More recently, patients who exhibit mild to severe CHD abnormalities have higher risks of CKD compared with controls without CHD^34^. Modeling kidney defects with *SMAD2* LoF mutations has been evolutionarily challenging because *Smad2* knockout mice are embryonically lethal, and the rescued mice have defective left–right patterning impacting multiple organs^43,44^. It remains unclear whether SMAD2 genetic variants linked to CHD can directly alter renal developmental programming and tissue patterning.

Given the substantial clinical associations between *SMAD2-*mediated CHD and CKD prevelence, we explored the roles of *SMAD2* LoF mutations in cell lineage specification and tissue patterning of the kidney. We postulated that defective renal cell and tissue patterning may occur earlier in development, such that mutations in *SMAD2* may result in abnormalities of the mesoderm and intermediate mesoderm (IM) progenitor cell lineages, which can ultimately disrupt patterning of the glomerulus and impair kidney function. We integrated human stem cell biology, CRISPR/Cas9 genome engineering, and microfluidic organs-on-chips systems to explore the questions from above.

## Results

### Loss of *SMAD2* posteriorizes mesoderm cells

During gastrulation, undifferentiated cells at the posterior edge of the epiblast give rise to the primitive streak, which undergoes an epithelial-mesenchymal transition to form mesoderm^1,45^. This mesoderm germ layer further specializes into distinct organ-specific precursors. Human iPS cells can be used to mimic these embryonic developmental processes by using appropriate signaling molecules that induce human iPS cell differentiation to a single mesodermal lineage. We generated isogenic iPS cells to create *SMAD2* LoF cell lines by non-homologous end joining (NHEJ) (**Extended Data Fig. 1**). To test whether *SMAD2* variants influence medial mesoderm derivation compared to wild-type (WT) cells, we differentiated both mutant and WT iPSCs into medial mesoderm cells using a 2-day differentiation protocol that activated TGFß and WNT signaling pathways using Activin A and glycogen synthase kinase-3ß (GSK-3ß) inhibition with CHIR99021, respectively (see **Methods**, **Fig. 1a**). Medial mesoderm serves as a common precursor for cardiogenic and nephrogenic transit amplifying cells^1^. Differentiated WT mesoderm cells demonstrated a distinct, tightly packed cobblestone morphology (**Fig. 1b**) and expressed medial mesoderm-specific markers (**Figs. 1c, d**). WT mesoderm consisted of a uniform expression of *Brachyury* (*T)* and *GSC* (*Goosecoid*) at both the mRNA and protein levels, a pattern consistent with the transient expression of these genes during gastrulation (**Figure 1d**). Thus, Activin A and GSK-3ß inhibition act in parallel to induce medial mesoderm formation in the WT cells.

**Figure 1:**
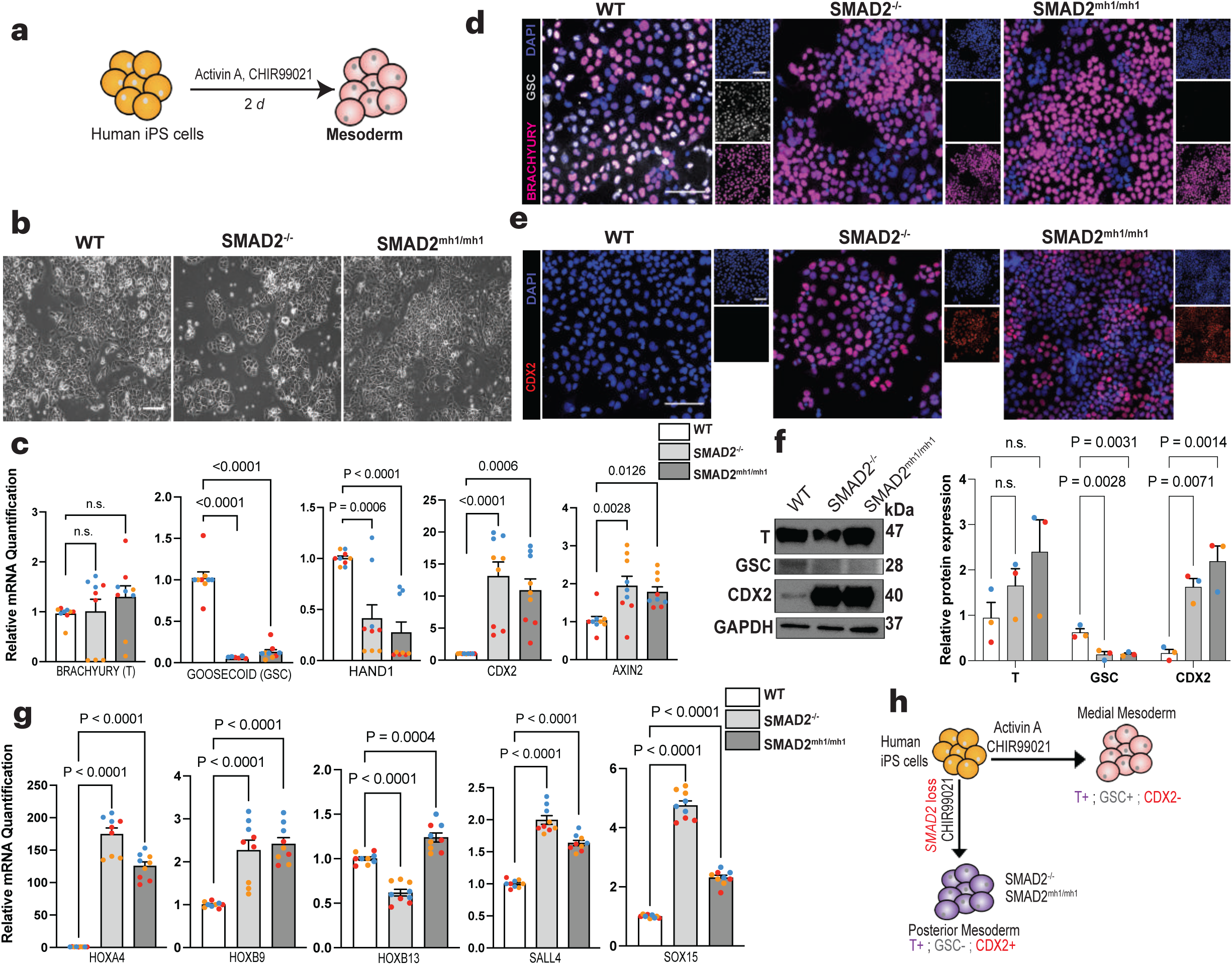
Loss of *SMAD2* shifts mesoderm lineage specification from medial to posterior fate. **a**, Schematic overview of the derivation of mesoderm cells from human iPS cells. **b**, Representative phase contrast images of iPS-derived mesoderm cells from WT and mutant lines. Scale bar, 275 µm. **c**, RNA expression of mesoderm lineage genes. Data are expressed relative to WT after normalizing to GAPDH and presented as mean values ± SEM (n = 3 independent experiments). *P*-value was calculated using one-way ANOVA with post hoc Dunnett’s test. **d**, Representative immunofluorescent images showing the expression of *Brachyury (T)* (magenta) and *Goosecoid* (*GSC*) (light gray) in the mesoderm cells. DAPI (blue) labels the cell nuclei. Scale bars, 100 µm. **e**, Representative immunofluorescent images showing the expression of *Caudal-type homeobox 2* (*CDX2*) (red), and DAPI (blue). **f**, Representative immunoblots on total cell lysates showing expression of lineage-specific markers. T expression in SMAD2^-/-^ was reduced compared to WT and *SMAD2^mh1/mh1^* mesoderm; GSC expression was obliterated in *SMAD2^-/-^* and *SMAD2^mh1^*^/*mh1*^ mesoderm; and significantly higher expression of *CDX2* was observed in *SMAD2*^-/-^ and *SMAD2^mh1^*^/*mh1*^ mesoderm. Data is representative of 3 biologically independent experiments. **g**, RNA expression of posterior mesoderm genes in the differentiated mesoderm cells. Expression of *Homeobox A4, B9, B13, Spalt Like Transcription Factor 4 (SALL4)*, and *SRY-Box Transcription Factor 15 (SOX15)* was significantly upregulated in *SMAD2*^-/-^ and *SMAD2^mh1^*^/*mh1*^ mesoderm cells compared to WT. Data are expressed relative to WT after normalizing to *GAPDH* as mean ± SEM (n = 3 independent experiments). *P*-value was calculated by one-way ANOVA with post hoc Dunnett’s test. **h**, Proposed model showing that stimulation of WT iPS cells with Activin A and CHIR99021 forms medial mesoderm cells positive for *T*, *GSC*, but negative for *CDX2*. However, in the absence of functional *SMAD2* protein, this differentiation is biased towards posterior mesoderm expressing only *CDX2* and *T* and negligible amount of GSC.

Transcriptional analysis of differentiated mutant mesoderm cells revealed a marked downregulation of *GSC* and *HAND1* mesoderm genes (**Fig. 1c**). Immunostaining confirmed that *SMAD2* mutant mesoderm cells, unlike WT cells, did not express GSC. Interestingly, in comparison to WT cells, *SMAD2* mutant cells expressed significantly higher levels of transcript and protein for a major determinant posterior mesoderm gene, *Caudal-type homeobox 2* (*CDX2*) (**Figs. 1c, e, f**). Expression of *CDX2* along with *T* in the mutant cell lines indicated a biased generation of posteriorized mesoderm^45^.

Prior experiments suggested that strong WNT pulse can induce late-stage primitive streak cells to give rise to posterior mesoderm cells^46^. We profiled the expression levels of the WNT target gene *AXIN2* in mutant mesoderm cells and noted significant upregulation, suggesting heightened canonical WNT activity in the mutants (**Fig. 1c**). We found that protein expression levels of *SMAD3* and *pSMAD3* increased in *SMAD2* mutant mesoderm cells compared to WT, indicating a compensatory role of *SMAD3* in the absence of *SMAD2* (**Extended Data Fig. 2a**). Noticeably, increased protein levels of *SMAD3* were not sufficient to specialize iPS cells toward medial mesoderm in *SMAD2* mutant lines (**Extended Data Fig. 2a**; **Figs. 1d, e**). RNA expression of posterior mesoderm genes *Homeobox A4, B9, B13*, *Spalt-Like Transcription Factor 4 (SALL4)*, and *SRY-Box Transcription Factor 15 (SOX15)* were significantly higher in *SMAD2* mutant mesoderm cells compared to WT, which confirmed our hypothesis that the absence of *SMAD2*/Activin A signaling induces a posterior mesodermal fate (**Fig. 1g**).

To better understand this altered mesoderm lineage determination, we asked whether loss of *SMAD2* can bias differentiation of mesoderm towards a more posterior fate due to the inability of the cells to respond to Activin A pulses. To test this, we differentiated WT and mutant iPS cell lines for 2 days with only CHIR99021 pulse (**Extended Data Fig. 2b**). Morphologically, the medial and posterior mesoderm cells were indistinguishable (**Fig. 1b** and **Extended Data** Fig. 2c). CHIR99021-pulsed WT and *SMAD2* mutant iPS cells robustly generated *T-* and *CDX2-*positive posterior mesoderm cells (**Extended Data Fig. 2d**). Our results echoed findings from embryo gastrulation, in which a strong WNT pulse induced *CDX2* positive mesoderm cells^47^. Next, we sought to characterize the mutant mesoderm cells at the molecular level. We observed no significant difference in expression of posterior mesoderm genes between the WT and the mutant cells (**Extended Data Fig. 2e**). Although GSK-3ß inhibition-mediated *ß-catenin* expression was enhanced in the presence of Activin A in the mutants, we observed no difference in mesoderm cells differentiated with CHIR99021 pulse alone (**Extended Data Fig. 2d**). We analyzed the cell culture supernatant for secreted Activin A morphogen when mesoderm was induced with CHIR99021 pulse but did not observe any signal (**Extended Data Fig. 2f**). Thus, we concluded that formation of the medial mesoderm was blocked by strong GSK-3ß inhibition in the mutants and the resulting mesoderm lineage was specified primarily via the WNT signaling pathway. Using iPS cell lines, we demonstrated the competence of *SMAD2* mutant mesoderm to adopt a posterior fate due to its inability to respond to Activin A signals. Our results provide genetic evidence that *SMAD2* functions as an important transcription factor, and its loss impairs and biases mesoderm lineage specification (**Fig. 1h**).

### *SMAD2* mutations exacerbate IM cell motility

In the developing embryo, most kidney cells, including glomerular podocytes, develop from the intermediate mesoderm (IM) cells, which, in turn, is derived from the mesoderm layer. To further understand the effects of *SMAD2* LoF variants on the developmental path to glomerular podocytes, we differentiated WT and mutant mesoderm cells towards nephrogenic IM cells via modulation of WNT and bone morphogenetic protein 7 (BMP7) signaling pathways (**Fig. 2a**). Wild-type IM cells exhibited a cobblestone-like morphology and were highly proliferative (**Fig. 2b**). In contrast, mutant cells failed to form the uniform cobblestone-like morphology (**Fig. 2b**), underwent cellular hypertrophy, developed a myofibroblast-like cytoarchitecture, and died during differentiation.

**Figure 2:**
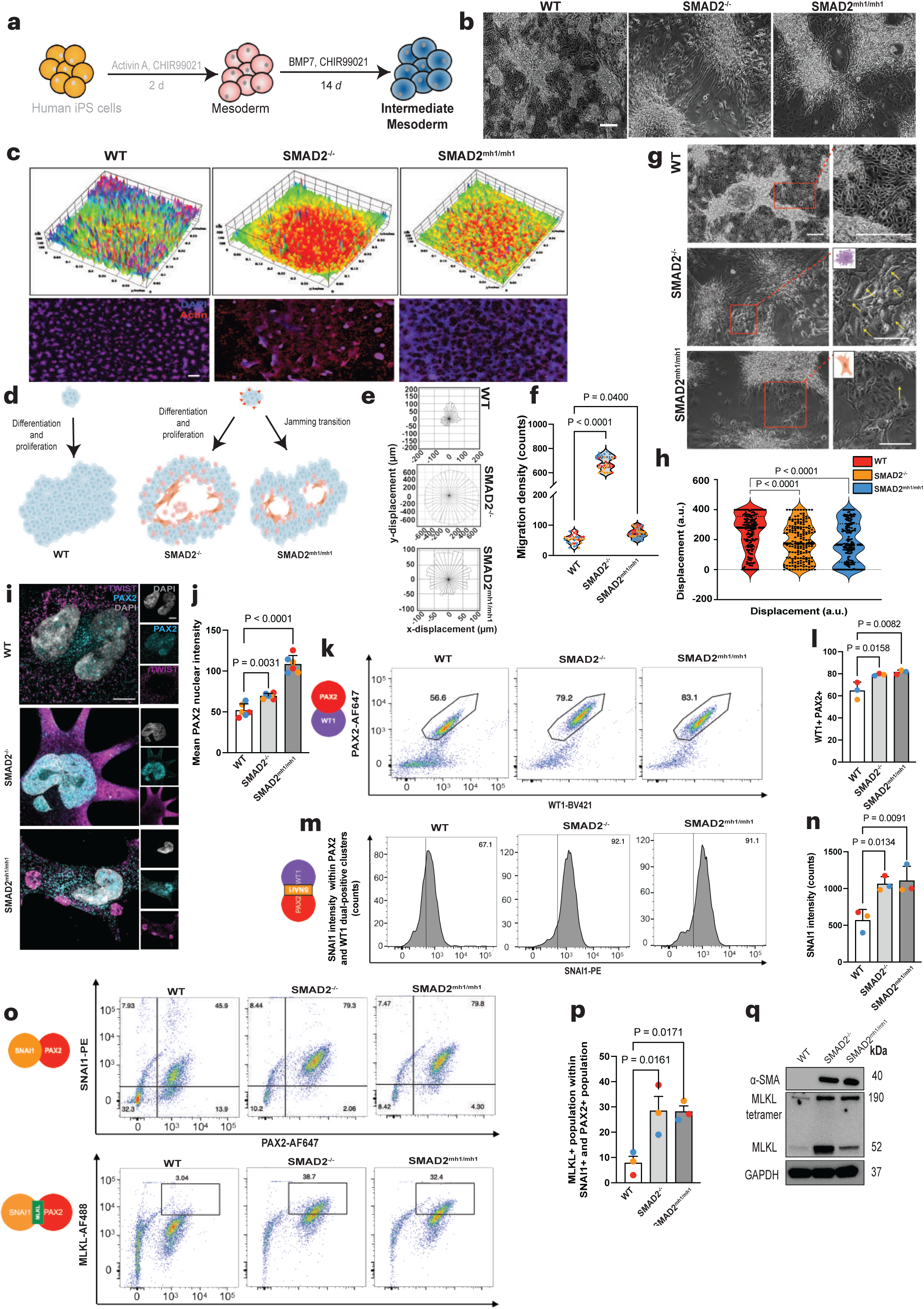
LoF mutations in the *SMAD2* gene directs aberrant differentiation of IM cells. **a**, Schematic representation of the derivation of IM cells from human iPS cell-derived mesoderm. **b**, Representative phase contrast images of WT, *SMAD2*^-/-^, and *SMAD2^mh1^*^/*mh1*^ IM cells. Scale bar, 275 µm. **c**, Representative 3-Dimensional (3D) surface plots of IM cells. Images (398 total) were captured per well from the center in a serpentine manner that represented 70% coverage of the well. These images were stitched with an in-built AI-based tool using the EVOS M7000 image analyses software. 3D surface plots were generated from the stitched images with grid size = 64 µm, scale = 0.82 µm steps, and z-scale = 0.38 µm steps. Immunofluorescence images of IM cells in the lower panel also represents 70% of the well. Cells were stained with DAPI (nuclei, blue) and Phalloidin (Actin, red). The WT IM cells showed a uniform growth pattern throughout the well, whereas the SMAD2^mh1/mh1^ IM had a distinct honeycomb with jammed cell clusters. SMAD2^-/-^ IM cells are highly mobile during differentiation with fewer jammed clusters. **d**, Schematic overview of IM cell differentiation pattern. *SMAD2*^-/-^ and *SMAD2^mh1/mh1^* IM displayed an EMT-like behavior with necroptotic blebs. *SMAD2^mh1/mh1^* IM formed a honeycomb network with jammed cell clusters. **e**, Representative rose plots of cell migration over a period of 167 h, and **f**. corresponding sector radii quantification from the rose plots. *P*-value was calculated by one-way ANOVA with post hoc Dunnett’s test. **g**, Representative phase contrast images of IM cells showing the morphology of cells localized between cell clusters. *SMAD2*^-/-^ IM cells between clusters formed numerous necroptotic blebs (indicated by yellow arrows) and exhibit myofibroblast-like phenotype. Conversely, *SMAD2^mh1/mh1^*IM cells that fail to join jammed clusters exhibit myofibroblast-like phenotype. Scale bars, 275 µm. **h**, Quantification of IM cell displacement along the XY Cartesian coordinates shows highest cell mobility in the *SMAD2*^-/-^ IM and constricted migration and jammed transition in the *SMAD2^mh1/mh1^*IM. *P*-value was calculated by one-way ANOVA with post hoc Dunnett’s test. **i**, Representative immunofluorescent images showing the expression of early nephrogenic IM lineage marker *Paired Box Homeotic Gene 2 (PAX2)* (cyan) and EMT maker *Twist Basic Helix-Loop-Helix Transcription Factor (TWIST)* (magenta). DAPI, (light grey) labels the nuclei, and the scale bars represent 5 µm. Single-channel images are shown next to the merged images for the indicated field of view. *TWIST* expression is pronounced and *PAX2* expression is mostly nuclear in the *SMAD2*^-/-^ and *SMAD2^mh1/mh1^* IM. Wild-type IM had a few *PAX2* nuclear puncta and diffused *TWIST* staining. **j**, Quantification of mean *PAX2* nuclear intensity. Data are mean, ± SD n = 5 cells. *P*-value was calculated by one-way ANOVA with post hoc Dunnett’s test. **k**, Representative flow cytometry plot showing co-expression of IM lineage markers AF647-*PAX2* and Wilms’ Tumor 1 (BV421-*WT1*). **l**, Dual positive cells were quantified as indicated in the gated populations. Data are mean ± SEM; n = 3 independent experiments; *P*-value was calculated by one-way ANOVA with post hoc Dunnett’s test. **m,** Representative flow cytometry of IM cell counts for EMT marker *Snail Family Zinc Finger 1* (PE-*SNAI1*) from cell populations that were double-positive for AF647-*PAX2* and BV421-WT1. **n**, SNAI1 intensity quantification. Data are mean ± SEM.; n = 3 independent experiments; p-value was calculated by one-way ANOVA with post hoc Dunnett’s test. **o**, Representative flow cytometry plot of IM cells gated for EMT marker PE-*SNAI1* and early nephrogenic IM maker AF647-*PAX2*. AF488-*MLKL* population was gated from PE-*SNAI1* and AF647-*PAX2* dual-positive populations. **p**, Quantification of AF488-*MLKL*-positive cells from PE-*SNAI1* and AF647-*PAX2* double-positive populations. Data are mean ± SEM; n = 3 independent experiments. *P*-value was calculated by paired t-test two-tailed. **q**, Representative immunoblots showing expression of myofibroblast marker *Alpha smooth muscle actin (αSMA)* in the differentiated IM. Cell lysates were probed for the presence of *MLKL* oligomers under non-denaturing, non-reducing conditions. Mutant IM cells expressed oligomerized form of *MLKL* centered at ∼190 kDa which indicated necroptosis-mediated cell death. Data is representative of 3 biologically independent experiments.

To further assess the abnormal morphology observed in the mutant cells exposed to IM induction conditions, we examined the cells by staining them with Phalloidin. We observed anisotropic remodeling of the cytoskeletal network with a higher signal of F-actin emanating from the periphery of the cell clusters in the mutants compared to WT (**Extended Data Fig. 3a**). These differentiating mutant cells underwent active rearrangement of the actin network, such that cells near the edges of the clusters extended long cytoplasmic protrusions to potentially help them move away from their original position and join the densely packed cell clusters (**Fig. 2b**). The mutant IM cells exhibited mesenchymal-like behavior with peripheral constriction of cell clusters, progressive changes in cell morphology, and a reduction in cell-to-cell contact. To visualize this differentiation pattern, we generated a 3D surface plot by fluorescence imaging of at least 70% of the cell culture surface area (2.45 cm^-2^ per well) using mosaic tiling (**Extended Data Fig. 3b, Fig. 2c**). The WT IM cells successfully differentiated and proliferated throughout the well and had a packed-phase. In contrast, the *SMAD2*^-/-^ IM cells mostly died and detached from the culture surface during differentiation as demonstrated by the red areas (empty spots in **Fig. 2c**) of the surface plot, forming only a few cell clusters (**Extended Data Fig. 3b**, **Figs. 2c-f**). It appears that the upregulation of *SMAD3* in the absence of *SMAD2* allowed these cells to secrete pathological matrices (including PAI-1 and Tenascin-C, **Extended Data Fig. 3m**) that provided strong adhesiveness to form densely packed clusters^48^ as evidenced in **Fig. 2c** and **Extended Data Figs. 3a, b**.

Strikingly, for the *SMAD2^mh1/mh1^* IM cells, we observed that differentiating IM cells migrated and formed smaller cell clusters but maintained a high degree of long-range orientational order (**Figs. 2b-f and Extended Data Figs. 3b, c**). The mutant cells that failed to form cell clusters appeared mesenchymal-like in morphology (with enlarged spindle shape), and developed bleb-or vesicle-like structures on their plasma membrane which was followed by cytoplasmic regurgitation and cell death (**Figs. 2c, g**, **Extended Data Figs. 3b, c**). Our 3D surface plot for *SMAD2^mh1/mh1^* IM also demonstrated cell clusters (green peaks) with long-range order and clear valleys (red spots) in between cell clusters (**Fig. 2c**). Time-lapse microscopy analysis confirmed that the physical location of differentiating groups of mutant IM cells was highly transitory, as these cells rapidly lost cell-cell contact and migrated away from their original position to join cell-dense clusters. The aberrant migratory patterns of the *SMAD2*^mh1/mh1^ IM cells seem to cause a loss in their ability to move, forming a jamming transition in which cells remained “caged” in the ‘jammed’ cluster (**Figs. 2e-g**, **Extended Data Figs. 3b**,). Although both the *SMAD2*^-/-^ and *SMAD2^mh1/mh1^* IM cells exhibited pronounced collective migration than the WT cells, the *SMAD2*^-/-^ underwent extensive cell-delamination, which counterbalanced overcrowding with fewer jamming transition (**Fig. 2e**, **Extended Data Fig. 3b**). To quantify these mesenchymal-like migratory phenotypes, we combined our time-lapse data with a semi-automated image analysis FIJI-pipeline and tracked the migration of cells. Rose plots of the data suggested that *SMAD2*^-/-^ IM cells were the most migratory, followed by *SMAD2^mh1/mh1^* and then WT IM cells (**Fig. 2e**, **Extended Data Fig. 3c**). The cell migration pattern appeared to be radial but spatially heterogeneous in magnitude, suggesting that the large-scale heterogeneities in the migration trajectories were dynamically initiated. As cells proliferated or increased in density within the jammed clusters, the adjoining mesenchymal-like cells between the clusters experienced restricted motion, underwent hypertrophy, adopted a myofibroblast-like morphology, or died (**Figs. 2f-h**).

### Mutant IM cells demonstrate an altered progenitor cell fate specification

BMP7 drives mesenchymal nephrogenic IM specification, whereas GSK-3ß inhibition primes mesenchymal-to-epithelial transition of IM cells towards multipotent nephron progenitor cells (NPCs)^49,50^. We examined whether BMP7 and WNT can instruct cell fate specification in WT and mutant isogenic cell lines towards nephrogenic IM. We performed high-resolution microscopy analysis on 14*d* IM cells for *PAX2* expression, which is an early molecular marker of IM cells. WT IM cells exhibited diffused nuclear expression of *PAX2* with a few punctae, whereas *SMAD2*^-/-^ and *SMAD2*^mh1/mh1^ IM cells had strong nuclear localization (**Figs. 2i, j**). As IM cells mature during development, BMP7 is known to significantly decrease the expression of PAX2^51^. However, high mean intensity of *PAX2* in the nucleus of the *SMAD2* mutant IM compared to WT cells highlighted the intriguing possibility of a partially-differentiated or an immature IM population. Expression of *SMAD3* can transcriptionally activate epithelial-mesenchymal transition (EMT) genes, including *Twist Basic Helix-Loop-Helix Transcription Factor* (*TWIST*) and *Snail Family Zinc Finger 1* (*SNAI1*)^52^. Therefore, we explored whether the differentiated IM cells expressed *TWIST*. High-resolution immunofluorescence studies revealed strong TWIST fluorescence in the mutant IM cells compared to the WT (**Fig. 2i**). Immunoblots also confirmed elevated expression of EMT makers (*TWIST, SNAI1*) in mutant IM cells (**Extended Data Fig. 3d**). Together, the expression of EMT markers (*TWIST, SNAI1*), acquisition of mesenchymal features, and motile phenotype in the mutants suggested a feedforward auto-regulatory loop that perturbed complete cell lineage commitment. Taken together, we observed a shift in cellular identity between WT and mutant cells undergoing IM differentiation, such that cell-cluster morphology, spatial interactions between cells, and the relative ratio of cytoplasmic/nuclear expression of early lineage-specific markers (e.g., *PAX2)* were all affected by the *SMAD2* mutations.

Intrigued by the dynamics in cell migration and aberrant expression of lineage markers, we sought to determine the cell phenotype between the cell clusters. WT IM cells grew to confluency within the tissue culture dish and exhibited a uniform cell distribution between cell clusters. In contrast, the *SMAD2* mutant cells developed aberrant differentiation patterns including jammed clusters, followed by cell death and myofibroblast-like cytoarchitecture (**Fig. 2g**). To further dissect the impact of WNT/BMP7 signaling on IM specification, we investigated cellular heterogeneity using multi-chromatic flow cytometry analysis. The frequency of cells that were dual positive for the IM markers *PAX2* and *WT1* was high in *SMAD2*^-/-^ (∼79.2%) and *SMAD2*^mh1/mh1^ (∼83.1%) compared to WT (∼56.6%) (**Figs. 2k, l**). This can result from the increased BMP7 signaling in the mutant cells, as evidenced by upregulated *pSMAD1* expression (**Extended Data Fig. 3e**). However, grouping for the *SNAI1*-positive population within the (*PAX2* and *WT1*) dual-positive IM clusters revealed significantly high *SNAI1* intensity in *SMAD2*^-/-^ (∼92.1%) and *SMAD2^mh1^*^/mh1^ (∼91.1%) compared to WT (∼67.1%) (**Figs. 2m, n**). Since BMP7 drives nephrogenic IM specification and helps in the expression of *PAX2* and *WT1*, we examined whether either of these transcription factors were partially responsible for the observed EMT phenotype in the mutant cells. We assayed for the percentage of cells that were dual positive for *SNAI1* (EMT) and *WT1* (IM) (**Extended Data Figs. 3f, g**), and *SNAI1* (EMT) and *PAX2* (IM) (**Fig. 2o**). For both conditions, we observed that compared to WT, mutant cells with upregulated expression of IM lineage markers were preferentially developing EMT-like phenotype.

### SMAD2 mutant IM cells are necroptotic

We noticed that mutant IM cells that fail to migrate towards jammed clusters underwent hypertrophy (**Fig. 2g**), developed plasmalemmal bubbles (**Fig. 2g**), and died (**Fig. 2g**, **Extended Data Figs. 3a, b**) which resembled necroptosis. We questioned whether *SMAD2* mutant IM cells in the EMT spectrum could undergo cell death by necroptosis. During necroptosis, *Mixed Lineage Kinase Domain Like Pseudokinase* (*MLKL*) forms oligomers that translocate to the plasma membrane and create pores. This phenomenon causes leakage of cytoplasmic contents that lead to cell death^53^. Analysis of the *MLKL*^+^ population within cell clusters dual positive for *SNAI1*(EMT) and *PAX2* (IM) (**Fig. 2o, p**) and *WT1* (IM)-positive populations (**Extended Data Figs. 3h, i**) revealed that mutant IM cells that have progressed towards the EMT spectrum die via *MLKL*-mediated necroptosis.

As MLKL oligomers are packaged with sensitive disulfide linkages, we ran our protein lysates under a non-reducing condition. Mutant IM cells expressed *MLKL* oligomers (∼190 kDa), which further confirmed our flow cytometry results (**Fig. 2q**). As differentiating mutant IM cells are more prone to cell death during the differentiation timeline, we characterized expression of the MLKL oligomers every alternate day for up to 14 days. We observed a drastic increase in MLKL expression for the mutant IM cells during differentiation compared to the WT, as evidenced by the bands centered at ∼190 kDa (oligomer) and 52 kDa (monomer) (**Extended Data Fig. 3j**); these results further validated our 3D surface plots (**Fig. 2c**) showing prominent loss of the differentiating mutant cell lines. Mutant IM cells also exhibited a gradual increase in expression of alpha smooth muscle actin (*αSMA*), indicating that these cells also develop myofibroblast-like features during lineage specification (**Fig. 2q and Extended Data Fig. 3k**). We then analyzed several canonical and non-canonical WNT ligands previously found to be associated with EMT and fibrosis^54^. Upregulated expression of WNT5A, WNT7A, WNT9A, and WNT10B revealed that mutant IM cells entered a highly mesenchymal state (**Extended Data Fig. 3l**). These mutant IM cells also displayed aberrant expression of mesenchymal/fibrotic markers, including Forkhead Box C2 (FOXC2), Forkhead Box L1 (FOXL1), TENASCIN C, and Plasminogen activator inhibitor-1 (PAI-1) (**Extended Data Fig. 3m**). While *pSMAD1* was previously shown to negatively regulate EMT and fibrosis,^55^ our results underscore the crucial balance between *SMAD* transcription factors during cell lineage specification, such that their aberrant expression nudges the developing cell lineage towards an EMT-fibrotic spectrum despite the upregulated BMP7 signaling.

### *SMAD2* mutations alter podocyte differentiation

To better understand the molecular underpinnings of *SMAD2*’s role in podocytogenesis, we differentiated IM cells towards a terminally differentiated podocyte fate by temporal control of WNT/BMP7/Activin A/RA pathways (**Fig. 3a**). Simultaneous activation of these pathways induces terminal nephrogenic differentiation *in utero*^56^ and *in vitro*^49^. Wild-type podocytes were characteristically large, arborized, and displayed primary and secondary foot processes, forming interdigitations between adjacent cells. Interestingly, the mutant podocytes displayed significantly high levels of cell death throughout the differentiation timeline, and the surviving cells were smaller in size and lacked foot processes, resembling an ‘effacement-like’ architecture (**Fig. 3b**, **Extended Data Fig. 4a**). Given the observed morphological abnormalities in the *SMAD2* mutant podocytes, we assessed lineage-specific marker expression in the differentiated cells using high-resolution microscopy. We observed high nephrin and podocin expression in the WT podocytes’ foot processes (**Fig. 3c**). Quantification of nephrin and podocin expression in the foot processes of the podocytes revealed significant enrichment of colocalized nephrin-podocin puncta in the foot processes as commonly observed in healthy podocytes (**Fig. 3d**) with high colocalization coefficients (**Fig. 3e**). On the contrary, the expression of nephrin and podocin were significantly downregulated in the mutant podocytes with a concomitant reduction in their overlap (**Figs. 3c-e**). The function of podocytes is enabled by their unique cytoarchitecture, particularly the development and stability of the foot processes, which possess highly ordered parallel contractile actin filaments. Synaptopodin plays an essential role in maintaining the stability of the foot processes^57^. Our WT podocytes demonstrated a web-like expression of Synaptopodin extending from the cytoplasm to the cell processes which was absent in the mutants (**Fig. 3c**). Quantification of immunofluorescence data further revealed differential subcellular protein localizations including decreased expression of nephrin, podocin, and synaptopodin in the cell body of the mutants compared to WT, (**Extended Data Fig. 4b**).

**Figure 3:**
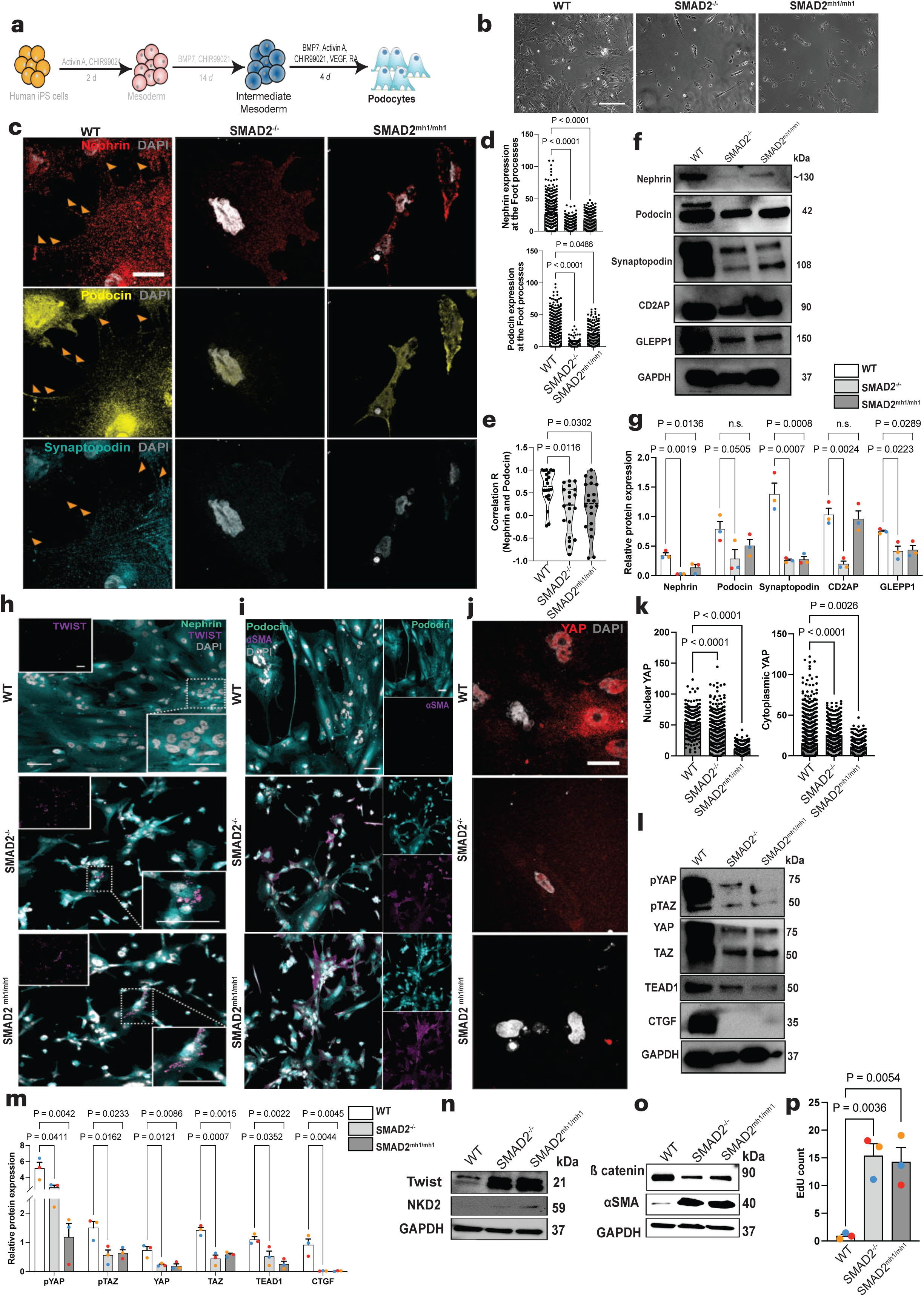
Mutant IM cells fail to specialize into terminally differentiated podocytes. **a**, Schematic overview of the derivation of podocytes from human iPS cell-derived IM. **b**, Representative phase contrast images of podocytes derived from WT and mutant IM cells. Scale bar, 275 µm. **c**, Representative immunofluorescent images showing the expression of podocyte lineage-specific markers: Nephrin(red), Podocin (yellow), Synaptopodin (cyan). The cell nuclei were counterstained with DAPI (grey). Orange arrows indicate punctate expression of nephrin and podocin. Scale bar, 100 µm for all images. **d**, Quantification of nephrin and podocin expression at the foot processes. *P*-value was calculated using one-way ANOVA analysis with post hoc Dunnett’s test. **e**, Pearson’s correlation coefficient for colocalized expression of nephrin and podocin expression at the foot processes. Data represent mean values ± sem; n = 3 independent experiments. *P*-value was calculated using one-way ANOVA analysis with post hoc Sidak’s test. **f**, Representative immunoblots showing expression levels of the podocyte markers nephrin, podocin, Synaptopodin, GLEPP1, CD2AP, and the housekeeping protein GAPDH. Data are representative of 3 biologically independent experiments. **g**, Relative protein expression of podocyte lineage markers across 3 biologically independent experiments. *P*-value was calculated using one-way ANOVA analysis with post hoc Dunnett’s test. **h**, Representative immunofluorescent images showing the expression of *Nephrin* (cyan) and EMT marker *TWIST* (magenta). Top left insets denote single-channel images of *TWIST* for the entire field of view. *TWIST* expression was pronounced in mutant podocytes. Bottom right insets denote zoomed-in regions marked by dashed white boxes, grey demarcates DAPI, cyan and magenta indicate *Nephrin* and *TWIST*, respectively. Scale bars, 100 µm. **i**, Representative immunofluorescent images showing the expression of *Podocin* (cyan) and *αSMA* (magenta) in podocytes. Single-channel images are shown next to the merged images for the entire field of view. *αSMA* expression is pronounced in mutant podocytes. Scale bar, 100 µm. **j**, Representative immunofluorescent images showing the expression of *YAP* (red) in differentiated podocytes. Scale bar, 100 µm for all images. **k**, Quantification of nuclear (active) and cytoplasmic *YAP* (inactive). *P*-value was calculated using one-way ANOVA analysis with post hoc Dunnett’s test. **l**, Representative immunoblots showing expression of *pYAP/pTAZ, YAP/TAZ* and its targets *TEAD1* and *CTGF*. Data are representative of 3 biologically independent experiments. **m**, Relative protein expression of *pYAP/pTAZ, YAP/TAZ* and its targets across 3 biologically independent experiments. *P*-value was calculated using one-way ANOVA analysis with post hoc Dunnett’s test. **n**, Representative immunoblots showing expression of EMT marker Twist and WNT antagonist *Naked Cuticle Homologue 2 (NKD2)*. n = 3 independent experiments. Their expressions were upregulated in the mutants. **o**, Representative immunoblots showing the expression of downstream WNT target *ß-catenin* and mesenchymal-myofibroblast marker *αSMA*. Reciprocal expression of *ß-catenin* and *αSMA* in the mutants indicated altered epithelization or mesenchymal phenotype of the podocytes. n = 3 independent experiments. **p**, Click-iT EdU quantification of podocytes displayed significantly high cell cycle activity in the mutant podocytes. Data represent mean values ± SD; n = 3 technical replicates. *P*-value was calculated using one-way ANOVA analysis with post hoc Dunnett’s test.

Apart from these lineage markers, we also examined glomerular epithelial protein-1 (*GLEPP1*) expression which is known to affect nephrin content in the foot processes^58,59^. We observed a significant downregulation of *GLEPP1* in the mutants (**Figs. 3f, g**). Several adaptor proteins, including *CD2AP*, are important for maintaining the foot process actin cytoskeleton. *CD2AP* binds both nephrin and podocin and links these proteins to actin^60^. We observed low expression of *CD2AP* in the *SMAD2*^-/-^ podocytes; however, we did not observe a significant difference in *CD2AP* localization in the *SMAD2^mh1^*^/*mh1*^ podocytes (**Fig. 3c**). While *CD2AP* expression was more spread out and punctate throughout the WT podocytes, its expression was more constricted near the nuclei for the mutants (**Extended Data Fig. 4c**). We also investigated the expression of *WT1* — an upstream regulator of many podocyte-specific makers^61^. WT podocytes had a nuclear expression of *WT1*, whereas its expression was either reduced or cytoplasmic for the mutant podocytes (**Extended Data Fig. 4c**).

Given these results, which established critical roles of BMP7/Activin A signaling towards podocyte differentiation from NPCs, we investigated protein level expression of key *SMAD* transcription factors. Western blot analysis revealed slightly higher expression of *pSMAD1* in the mutant podocytes, indicating mildly upregulated BMP7 signaling, and significantly high *SMAD3* which contributes to a mesenchymal phenotype (**Extended Data Figs. 4d, e**). *SMAD2* was strongly expressed in the WT cells whereas in the mutants, its expression was completely obliterated and accompanied by upregulation of *SMAD3* (**Extended Data Figs. 4c, d**). We also profiled for the expression of inhibitory SMADs including *SMAD6* and *SMAD7*. *SMAD6* and *SMAD7* are known to negatively regulate SMAD3 activation and functions^62^ and reduction of *SMAD7* promotes kidney fibrosis in a *SMAD3*-dependent manner^63^. Previous studies have shown that *SMAD7* deficiency leads to progressive fibrosis, inflammation in a unilateral ureteral obstruction (UUO) model of renal fibrosis^63,64^. Downregulation of the inhibitory SMADs in the mutant podocytes indicated an EMT-like phenotype in the mutants^65^ (**Extended Data Figs. 4d, e**).

### *SMAD2* mutant podocytes acquire mesenchymal phenotype

Next, we assessed whether podocytes generated from the mutant cell lines might also display mesenchymal properties. Immunofluorescence studies revealed punctate expression of *TWIST* in the mutant podocytes (**Fig. 3h**). *TWIST* is a potent EMT inducer, and its expression upon epithelial differentiation indicates incomplete epithelization of NPCs^66^. *TWIST* was downregulated in the WT podocytes, indicating that its suppression is crucial for maintaining the epithelial cell state (**Fig. 3h**). *SMAD3* can be a potent inducer of a mesenchymal gene expression program and enable transition of epithelial cells into *αSMA*-expressing myofibroblasts^52^. Similarly, activation of this EMT program can inhibit the terminal differentiation of podocytes. Our immunofluorescence studies corroborated loss of apicobasal polarity of the mutant podocytes, with strong *αSMA* emanating from the intercellular junction between adjacent podocytes suggesting a myofibroblast-like cell state (**Fig. 3i**). We concluded that a dysregulated *TGF-ß* axis with sustained difference in *SMAD2* and *SMAD3* expression in the mutant cells results in reduction of epithelial markers and induces evasive survival mechanisms (initiation of an EMT program) due to inability of the differentiating mutant cells to respond to SMAD2/Activin A signals.

### *SMAD2* mutant podocytes are developmentally immature

EMT is reciprocally linked to epithelization of NPCs toward terminal differentiation of podocytes^66^ and YAP/TAZ plays an important role in maintaining the terminally-differentiated podocyte fate. Nuclear-localized YAP is essential for podocyte viability^67–69^ and podocyte-specific loss of *YAP* leads to glomerulosclerosis^70^. Given the close association between *YAP/TAZ* and TGF-ß pathways, we probed for the expression of YAP and its molecular targets in the podocytes. We observed significant downregulation of nuclear *YAP* in the mutant podocytes (**Figs. 3j, k**). We also found a significant downregulation of *pYAP/pTAZ* and its targets *TEAD1* and *CTGF* in the mutant podocytes (**Figs. 3l, m**). In the nucleus, *YAP* binds to *TEAD* and mediates the expression of *CTGF*, which modulates cell-matrix interactions^71^. We observed an almost complete loss of *CTGF* from the mutant podocytes (**Figs. 3l, m**). Since *YAP* is involved in podocyte terminal differentiation, we investigated the expression of *PAX2* (early IM marker) and *WT1* (upstream regulator of podocyte-specific markers)^72^. We observed higher and lower nuclear expression of *PAX2* and *WT1* in the mutants respectively, which was completely reversed in the WT (**Extended Data Figs. 4 c, f**). *YAP* also plays an essential role in F-actin assembly^73–75^ and prevents the pro-apoptotic signaling of podocytes during injury^76,77^. While our WT podocytes displayed an outward distribution of actin which is critical for podocyte architecture and function (**Extended Data Fig. 4g**), the distribution of F-actin was lost in the mutants, with peripheral folding of actin bundles indicating effacement. *Synaptopodin* also associates with F-actin and helps maintain podocyte architecture ^57^. However, *Synaptopodin* expression was downregulated in the mutants (**Figs. 3c, e, f**). Cell viability was also significantly lower in the mutants (**Extended Data Fig. 4g**), likely due to defects in cell structure as suggested by actin disassembly (**Extended Data Fig. 4h**) and defective foot processes^68^.

Reduced expression of podocyte lineage specific markers (*nephrin*, *podocin*, and *synaptopodin*), coupled with upregulated expression of the *TWIST* (an EMT and stem-like signature)^78^ indicate stemness or less specialized cell state in the mutant podocytes (**Figs. 3h, n**). A previous study identified Naked Cuticle Homologue 2 (*NKD2*) as a fibrogenic biomarker in the human kidneys^79^. Expression of the WNT antagonist *NKD2* (**Fig. 3n**), combined with the observed EMT program and reciprocal expression of *ß-catenin* (low) and *αSMA* (high) (**Fig. 3o**) indicated incomplete epithelization and a fibrotic phenotype in the mutant podocytes. Analyses of previously published data on healthy and diseased human kidney podocytes revealed that CKD-injured podocytes expressed *TWIST1* and both CKD and AKI-injured podocytes expressed *TWIST* and *NKD2* when compared to healthy podocytes (**Extended Data Fig. 4i**).Injured podocytes can re-enter the cell cycle, but instead of progressing towards mitosis, the cell cycle is often arrested at the G1 or G2/M restriction point^80^. Given the levels of EMT and fibrogenic factors observed, we examined whether the mutant podocytes were terminally differentiated. We administered pulses of 5-ethynyl-2-deoxyuridine (EdU) after podocyte differentiation for 24 h. Our results showed enhanced EdU incorporation in the mutants relative to WT, indicating a lack of terminal differentiation (**Fig. 3p**). Taken together, the combined activation of an EMT program (*TWIST*) and expression of WNT antagonist (*NKD2*) and myofibroblast marker (*αSMA*) and downregulation of ß catenin highlight the failure of podocyte epithelization in the mutant cells compared to WT. The EMT program results in survival of some mutant cells, but the resulting cells are unable to develop mature podocyte phenotype or highly specialized foot processes required for proper organ function. We wondered if reducing *SMAD3* expression could rescue podocyte viability and cytoarchitecture. However, the mutant podocytes after *SMAD3* knockdown demonstrated excessive cell rounding, effacement of foot processes (indicated with orange arrows in **Extended Data Fig. 4j**), and cell death (**Extended Data Fig. 4j**) with a concomitant reduction in cell size (**Extended Data Fig. 4k)**, cell viability (**Extended Data Fig. 4l)** and reduction in *Synaptopodin* expression (**Extended Data Fig. 4m)**. Our results show the importance of SMAD2/Activin signaling in podocytogenesis.

### Human iPS cell-derived endothelial cells carrying SMAD2 mutations develop aberrant mesenchymal-like characteristics

Endothelial cells form an important component of the glomerular filtration barrier (GFB), where they interface with podocytes and the GBM to enable the kidney’s blood filtration function. To engineer a model of the GFB, we also differentiated isogenic endothelial cells from the WT and mutant iPS cells using our optimized protocol^81,82^ (**Fig. 4a**). Differentiated mutant and WT cells were MAC-sorted via CD31^+^ and CD144^+^ expression, followed by propagation with temporal exposure to conditioned media (see **Methods**) with VEGF-A supplementation for an additional four days. Differentiated endothelial cells (ECs) demonstrated a packed elongated morphology (**Fig. 4b**). The differentiation yield of mutant ECs (<20%) was significantly reduced compared to WT (∼80%) (**Fig. 4c**), indicating that abrogation of SMAD2 also attenuates EC differentiation. Immunofluorescence analysis revealed that the sorted WT and mutant ECs expressed lineage-specific markers, including platelet endothelial cell adhesion molecule (*PECAM1*/CD31) and Vascular Endothelial-Cadherin (*VECAD*) (**Fig. 4d**). Since we observed EMT-fibrotic behavior in differentiating IM cells and podocytes, we investigated transcript level expression of endothelial-mesenchymal (EndoMT) markers in our iPS-derived ECs. Intriguingly, we found significant upregulation of Intercellular adhesion molecules 1 (*ICAM-1*)/CD54, *smoothelin*, and *vimentin* in the mutant ECs (**Extended Data Fig. 5a)**. The mutant ECs cultured in maintenance media for two weeks underwent hypertrophy, adopted a Transgelin-positive fibroblastoid morphology, and expressed *TWIST* (**Fig. 4f** and **Extended Data Fig. 5b**). These cells also underwent junctional remodeling, as evidenced by the loss of *PECAM1*, and expressed late EndoMT marker *ICAM1*/CD54 (**Fig. 4e**). Together, these results demonstrate that defective cellular differentiation due to LoF *SMAD2* mutations significantly impact the cell lineage determination of both glomerular epithelium (podocytes) and vascular endothelium at the cellular level. Next, we examined the impact of the LoF *SMAD2* mutations on the formation and structure of the epithelial-endothelial tissue interface as described below.

**Figure 4:**
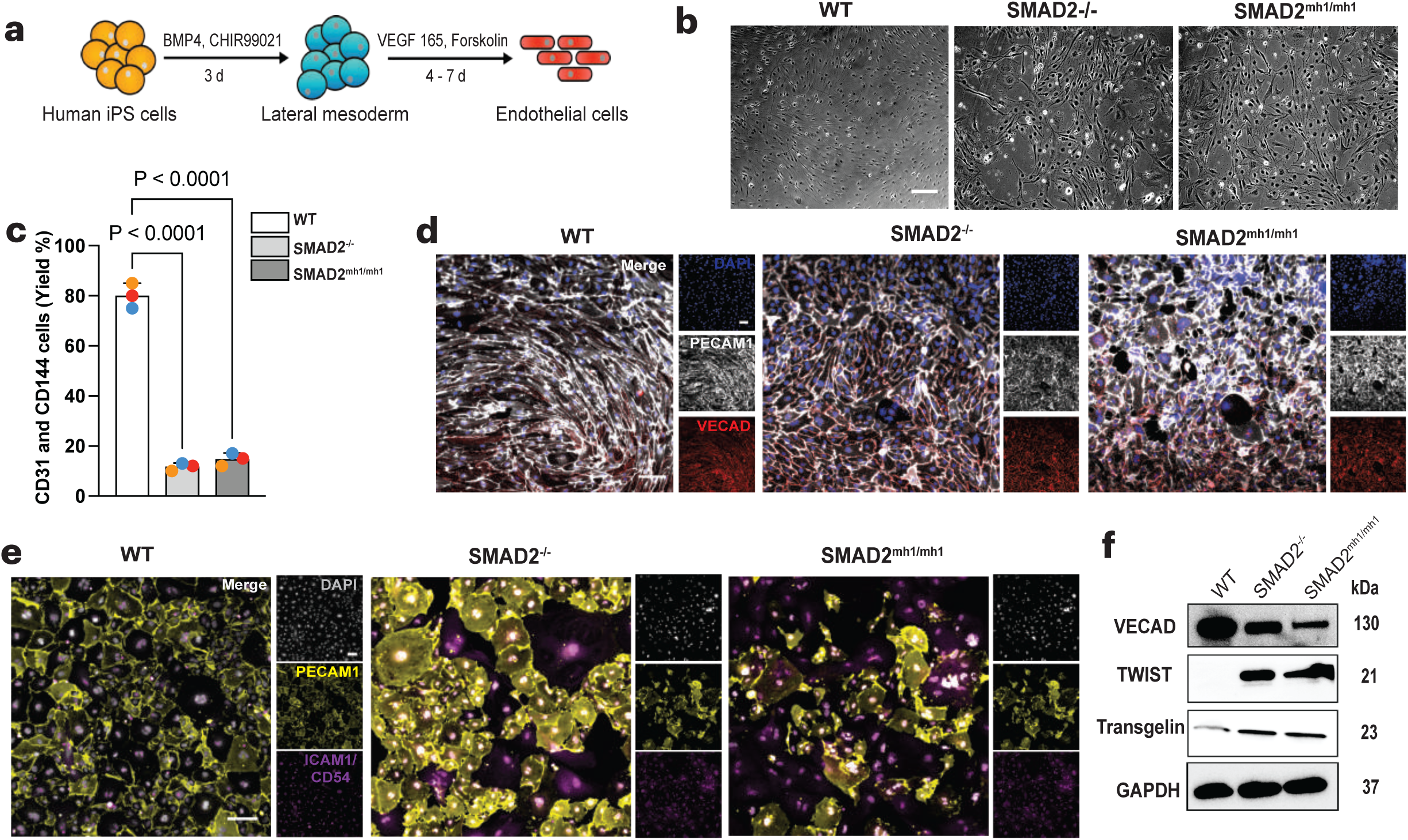
Mesenchymal phenotype observed in vascular cells differentiated from *SMAD2* mutant iPS cells. **a**, Schematic representation of the derivation of endothelial cells. **b**, Representative phase contrast images of the differentiated endothelial cells two additional culture days after MAC sorting. Scale bar, 275 µm. **c**, Endothelial cell differentiation yield calculated after MAC sorting. *SMAD2*^-/-^ and *SMAD2^mh1/mh1^* endothelial cells had a lower differentiation yield compared to the WT. Data are mean values ± SEM; n = 3 independent experiments. *P*-value was calculated by one-way ANOVA with post hoc Dunnett’s test. **d**, Representative immunofluorescent images showing the expression of endothelial marker *Platelet endothelial cell adhesion molecule (PECAM1*/CD31, light grey) and *Vascular Endothelial-Cadherin (VECAD*, red). Single-channel images are shown next to the merged images for the entire field of view. DAPI, shown in blue, labels the nuclei, and the scale bar represent 100 µm. **e**, Representative immunofluorescent images showing the expression of endothelial marker *PECAM1* (yellow) and mesenchymal marker *Intercellular Adhesion Molecule 1 (ICAM1/CD54)* in WT, *SMAD2*^-/-^, and *SMAD2^mh1/mh1^* endothelial cells 2 weeks after differentiation. Single-channel images are shown next to the merged images for the entire field of view. DAPI, shown in light grey, labels the nuclei, and the scale bar represent 100 µm for all images. **f**, Representative immunoblots showing the expression of EC, EMT, and mesenchymal markers in the mutant endothelial cells, including VECAD, *TWIST*, and *Transgelin*. Data is representative of 2 biologically independent experiments.

### Podocyte-endothelium tissue interfaces from LoF *SMAD2* mutant iPS cells fail to form a functional glomerular filtration barrier

During the S-shaped body stage of human glomerulogenesis, an endothelium-lined capillary invades the glomerular cleft^83^. The resulting topographic tissue arrangement allows NPC cells to acquire discrete regional identities as a function of their position. As the glomerulus matures, ECs contribute to podocyte development and cell lineage specification from immature columnar epithelial cells with adherens junctions into specialized epithelial cells (podocytes) with interdigitating foot processes . The developing podocytes wrap around the glomerular capillaries and interact with each other through a specialized type of cell junction called the slit diaphragm, which is located between interdigitating foot processes (**Fig. 5a**)^86^. Bioengineered microdevices like organ-on-a-chip systems can recapitulate key aspects of glomerular structure and function. To model glomerular tissue development process *in vitro*, we differentiated podocytes from IM cells and interfaced them with isogenic ECs in a microfluidic chip (**Fig. 5b**, **Extended Data Figs. 6a, b**). Mutant and WT IM cells were differentiated (in separate chips for each cell line) into podocytes in the top channel while the differentiated ECs were maintained in the bottom channel using their respective cell culture media (**Extended Data Fig. 6c**). After podocyte differentiation and EC maintenance, the podocyte and EC layers were confluent in the WT chips. However, the mutant podocytes and endothelial cells lost the characteristic expression and pattern of their respective lineage-specific markers and appeared less confluent. (**Extended Data Figs. 6c, d**). Interestingly, for the *SMAD2*^-/-^ chips, immunofluorescence studies revealed that the differentiation efficiency and morphology of the podocytes were compromised, such that only the cells in the chip areas interfacing with isogenic ECs (interface section of **Extended Data Fig. 6e**) had improved morphology. When podocytes were not in proximity (or interfaced) with the ECs, the mutant cells mostly died during differentiation (apical side in **Extended Data Figs. 6e, f**). The WT chips had uniform epithelial layers consisting of nephrin^+^ podocytes; for the mutant chips, the podocyte layers were discontinuous (**Extended Data Fig. 6c**. In the WT chips, ECs expressed cell junction marker *PECAM1*, and the mutant chips displayed diffused *PECAM1* expression throughout the cell cytoplasm (**Fig. 5c**). For the mutant microfluidic chips, reduced extension of podocyte foot processes and altered expression of lineage markers indicated an ‘effacement-like’ cell architecture (**Figs. 5c-e**). Prolonged maintenance of the mutant chips demonstrated extensive cell delamination (**Figs. 5c-e**). Our filtration assay revealed exacerbating proteinuria (pathologic leakage of albumin) for the mutant microfluidic chips (**Fig. 5f**). Together, these results underscore the impact of pathogenic mutations on glomerular tissue development, podocyte-endothelium cross-talk, and organ function, providing experimental evidence for kidney malformation and dysfunction clinically observed in patients with CHD^87–89^.

**Figure 5:**
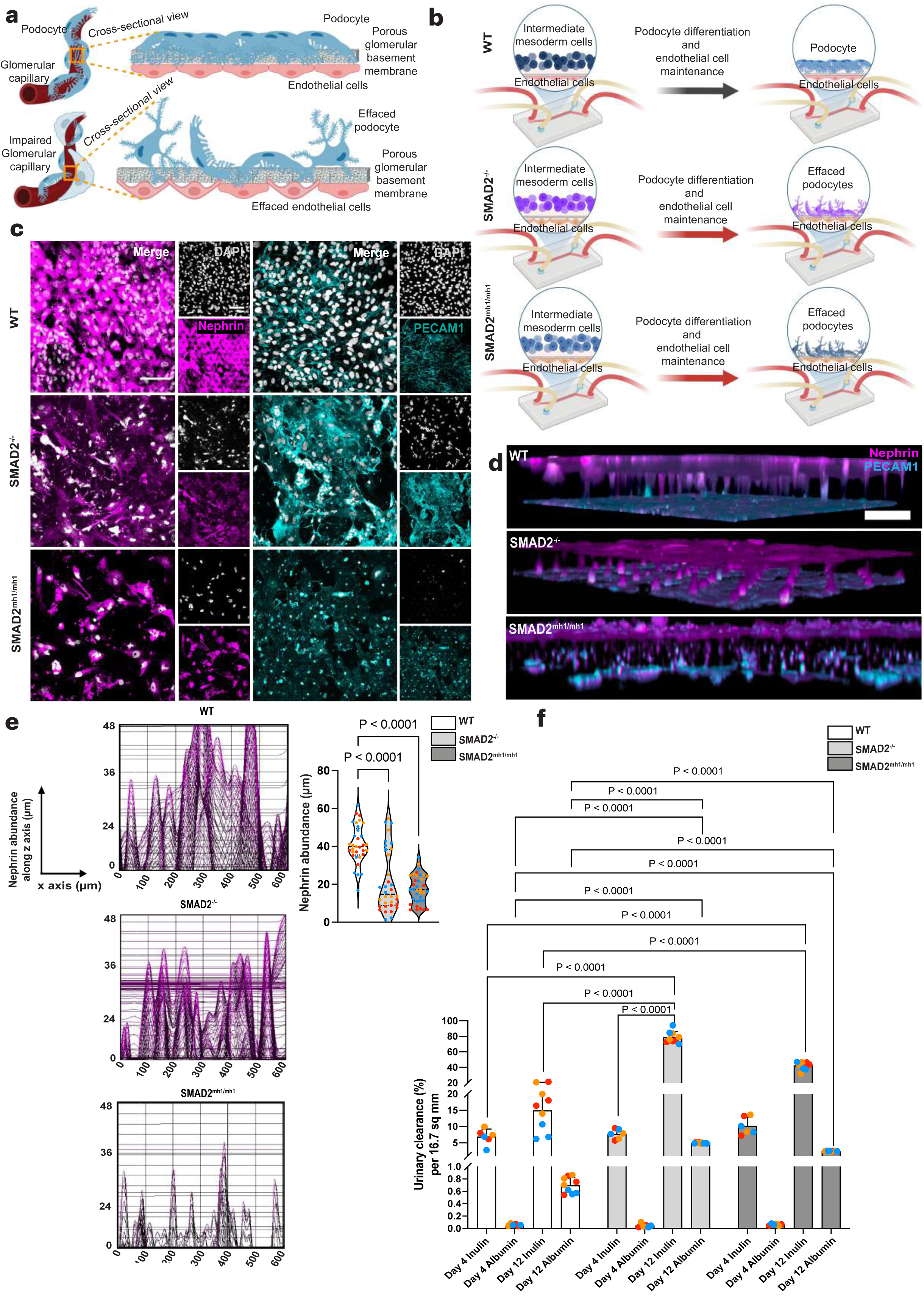
Engineering isogenic glomerular filtration barrier using microfluidic organs-on-chips devices. **a**, Schematic representation of normal and diseased/damaged glomerular capillary wall. **b**, Schematic representation of isogenic glomerular filtration barrier. The iPS-derived podocytes and endothelial cells were separated by a porous PDMS membrane. Under healthy conditions, podocytes wrap around the glomerular capillaries and allow filtration of molecules from blood. However, the mutant podocytes rapidly lose their specialized architecture followed by effacement and detachment. This phenomenon exposes the capillary towards the epithelial (urinary) compartment, leading to albuminuria. **c**, Representative immunofluorescence images of WT, *SMAD2*^-/-^, and *SMAD2^mh1/mh1^* iPS cell-derived podocytes (*Nephrin*) and endothelial cells (*PECAM1*) differentiated in the Glomerulus-on-a-chip platform. Scale bar, 100 µm. **d**, Cross-sectional views of the chips immunostained for *Nephrin* (podocytes, magenta, urinary channel) and *PECAM1* (endothelial cells, cyan, capillary channel). Scale bar, 30 µm. **e**, Nephrin abundance along the z axes of the chips. **f**, Quantification of the urinary clearance of albumin and inulin continuously perfused for 8 h into the capillary channel of the glomerulus-on-a-chip platform that was lined by isogenic podocytes and endothelial cells. Data are expressed as mean values ± SD; n = 3 independent experiments. *P*-values were calculated by two-way ANOVA with post hoc Tukey’s multiple comparisons test.

## Discussion

The presence of LoF mutations in genes associated with organ development and patterning, including *SMAD2*, can lead to anatomical defects and fetal mortality^90^. We sought to explore the impact of CHD-associated *SMAD2* LoF mutations on podocytogenesis and development of the glomerular filtration barrier, the primary site for blood filtration in the kidneys.

To study aberrant differentiation towards kidney podocytes, we introduced LoF *SMAD2* variants into human iPS cells. We employed a method for directed differentiation towards major cell lineages and delineated molecular mechanisms involved in cell fate specifications. The absence of functional *SMAD2* resulted in a biased mesoderm lineage commitment towards posterior mesoderm, and the mutant mesoderm cells remained largely nonresponsive to Activin A pulses. During mesoderm patterning, Brachyury (T) interacts with *SMAD2/3* to regulate the expression of mesoderm lineage-specific genes *GSC* and *HAND1*. *GSC* expression depends on a synergistic interaction between XTWN and Homeobox MIXER transcription factors induced by canonical WNT signaling and Activin-like factors through SMAD2, respectively^91,92^. In vivo, *HAND1* gives rise to the heart tube, the primary precursor of the left ventricular chamber and, to a smaller extent, the atria^93^. We note that loss of HAND1 and GSC in the mutants can be linked to the role of SMAD2 LoF mutations in CHD and observed clinical heart anomalies, including left-right asymmetry, abnormal heart looping, and tetralogy of Fallot which can impact kidney function.

In this study, mutant mesoderm cells failed to specialize towards definitive IM cells, as evidenced by the expression of TWIST, SNAI1, *α*SMA, MLKL, despite temporal stimulation with exogenous WNT and BMP7. Mutant IM cells progressed toward an EMT-like spectrum owing to elevated BMP7 and reduced WNT signaling. Upregulated *pSMAD1* and *SMAD3* expression may explain the preferential sensitivity of the mutant IM cells for acquiring EMT-like states with simultaneous dysregulation of *PAX2* expression. Higher nuclear localization of *PAX2* coupled with low *WT1* expression in the mutants could indicate a phenotype associated with severe glomerulopathy^94,95^ or nephronophthisis^96^. Differentiated mutant IM cells failed to specialize into arborized podocytes, and the development of an EMT-like state indicated an immature cell fate. The mutant podocytes, which expressed lineage-specific markers, were largely immature and failed to develop phenotypic characteristics such as interdigitated foot processes. Our findings suggest that elevated levels of *pSMAD1* (BMP7 signaling), *SMAD3*, and *NKD2* (a Wnt antagonist and inhibitor of epithelization) could be a means by which mutant IM cells counterbalance WNT signaling (reduced *ß-catenin* expression) in differentiating podocytes and drive the expression of *TWIST* and *αSMA*.

Intriguingly, when compared to WT, the mutant podocytes initially did not show significant phenotypic and functional difference when differentiated while interfaced with ECs in the microfluidic kidney glomerulus-on-a-chip. This is likely due to paracrine signals at the podocyte— endothelium interface, supporting the formation of primitive tissue structure. However, the engineered GFB deteriorated over time for the mutant cells as the chips were maintained for prolonged periods of time (7 or more days after differentiation). Specifically, extended culture of the glomerulus-on-a-chip resulted in delaminated patches of the mutant epithelium whereas the ECs on the basal side of the tissue model lost expression of their characteristic lineage-specific markers and acquired a fibroblastoid morphology. Selective molecular filtration analysis revealed that the aberrant tissue morphologies observed in the mutant microphysiological models resulted in exacerbated proteinuria as commonly observed in patients with CHD^11,16,31–39,89,97,98^.

In conclusion, we integrated CRISPR-based disease modeling with stem cell and microfluidic organ-on-a-chip technologies to systematically interrogate the impact of LoF *SMAD2* genetic variants on nephrogenic cell lineage specification and GFB function. We developed isogenic glomerulus-on-a-chip systems that recapitulated key aspects of GFB development and function with precise isogenic comparisons of LoF *SMAD2* variants. This strategy enabled the discovery of previously unexplored roles of LoF *SMAD2* variants on defective kidney tissue patterning and glomerular dysfunction. Studying pathogenic variants in the *SMAD2* gene has been evolutionary challenging due to the lack of human-specific models. *Smad2* knockout mice have defective organ pattering and are embryonically lethal whereas rescued mice develop holoprosencephaly and cyclopia^43,44,99^. Conditional *smad2* knockout mice develop *SMAD3*-mediated tubulointerstitial fibrosis which could be rescued after *Smad2* overexpression^52^. Patients undergoing peritoneal dialysis have increased TGF-β1 levels in peritoneal fluid, and the patients develop peritoneal fibrosis mediated by *SMAD3* but not *SMAD2*^100^, whereas *SMAD3*-deficient mice are protected from peritoneal fibrosis^100^. Despite the rising number of genetic variants linked to CHD and the extensive clinical heterogeneity among fetuses with *SMAD2* mutations, complex genetic disorders remain substantially challenging to model and treat effectively. Our results underscore the utility of genome-engineered human stem cell-based models and organ-on-a-chip technology to illuminate the cellular and molecular mechanisms of congenital kidney defects arising from mutations associated with extra-renal complications and multi-organ failure. We advocate for routine molecular diagnostic screening of kidney function in patients with complex CHD arising from genetic mutations, including *SMAD2,* because GFR reduction in CHD is strongly associated with mortality^16^. Our study also provides a viable microphysiological platform for future therapeutic discovery.

## Materials and Methods

### Generation of *SMAD2* mutant lines

Isogenic PGP1 iPS cell lines carrying *SMAD2* LoF mutations were generated using CRISPR/Cas9 technology. *SMAD2* LoF iPS cells were generated by non-homologous end joining (NHEJ) as previously described^101^ Two micrograms of plasmid-expressing Cas9 (PX459v2 from Addgene) were co-transfected with 2 µg plasmid-expressing guide RNA using a stem cell nucleofector kit (Lonza, Amaxa). Multiple guide RNAs were used to generate a collection of isogenic mutant cell lines. Gene edited clones were selected using puromycin (Gibco), and then expanded in culture followed by genotyping as previously described^101^. Each cell line was cultured for more than 8 passages prior to use before experiments.

Gene-edited cells were subcloned twice and validated by Sanger sequencing. Subcloning involved dissociation of iPS cells by pipetting and filtering through a 60-µm cell strainer. Cells were then plated onto a 6-well plate using mTeSR1 medium (StemCell Technologies; 85870) and Y27632 Rock inhibitor (10 nM) (R&D systems, 1254). Individual colonies were allowed to grow to about 300 cells, and clones were picked and separately placed into individual wells of a 96-well plate. Once the replated clones grew to approximately 85% confluency, iPS cells were collected and processed for PCR, where the sequence of the variant was amplified. To ensure purity of iPS cell clones, the cells were then subjected to an additional set of sub-cloning, and PCR amplified fragments were submitted for Sanger sequencing and next generation sequencing. Confirmation of sequencing was performed using bioinformatic analysis, DNA-star software (Lasergene 17), and Integrated Genomics Viewer (IGV). Two independent clones were created for each LoF *SMAD2* genotype (**Extended Data Figs. 1a-c**). *SMAD2* was stably expressed in the WT, but its expression was abrogated in the mutant lines (**Extended Data Fig. 1d**). All cell lines grew as smooth-edged, tightly packed colonies and were karyotypically normal (**Extended Data Figs. 1e, f**).

### Cell Culture

All cell lines used for this study were obtained under appropriate material transfer agreements and approved by all involved institutional review boards. All cells were tested for and shown to be free of mycoplasma contamination (Mycoplasma PCR Detection Kit from abm, G238). Human (iPS) cell line -PGP-1 (the Personal Genome Project-1) used for this study were propagated on tissue culture plates coated with Matrigel (VWR; 75796-278) by using mTeSR1 (StemCell Technologies; 85870) medium without antibiotics. The cells were split (at 1:6) every 4 to 5 days by treatment with Accutase (Thermo/Life Technologies; A1110501). The minimize cell loss, the mutant lines were split (1:3) every 4 to 5 days. All cells were propagated in a 37 °C incubator with 5% CO_2_.

### Differentiation of mesoderm, IM, and podocytes

Human iPS cells were dislodged with enzyme-free dissociation buffer (Gibco, 13150-016) and centrifuged twice at 200 g for 5 min each in advanced DMEM/F12 (Gibco; 12634010) to remove residual Matrigel or stem cell culture media components. The supernatant was aspirated, and the cell pellet was resuspended in mesoderm induction media consisting of DMEM/F12 with GlutaMax (GIBCO, 10565018) supplemented with 100 ng/ml activin A (Thermo Fisher Scientific, PHC9561), 3 μM CHIR99021 (Stemgent, 04-0004), 10 μM Y27632 (TOCRIS, 1254), and 1× B27 serum-free supplement (GIBCO, 17504044). Cells were plated on laminin-511-E8-coated plates (Takara; T-304) at a seeding density of 150,000 cells per well of a 12-well plate. The cells were cultured for 2 days with daily media change, and after two days, intermediate mesoderm differentiation (IM) was induced by feeding the cells with IM induction media consisting of DMEM/F12 with GlutaMax supplemented with 100 ng/ml BMP7 (Thermo Fisher Scientific, PHC9541), 3 μM CHIR99021 (Stemgent, 04-0004), and 1× B27 serum-free supplement (GIBCO, 17504044) for 14 days. For podocyte induction, IM cells were trypsinized with 0.25% trypsin-EDTA (Gibco; 25200-056), individualized by repeated pipetting, and plated on freshly prepared laminin-511-E8-coated plates (Takara; T-304) at a seeding density of 150,000 cells per well on a 12-well plate. The cells were fed for 4 days with podocyte induction media consisting of DMEM/F12 with GlutaMax supplemented with 100 ng/ml BMP7, 100 ng/ml activin A, 50 ng/ml VEGF (Thermo Fisher Scientific, PHC9391), 3 μM CHIR99021, 1× B27 serum-free supplement, and 0.1 μM all-trans retinoic acid (Stem Cell Technologies, 72262).

### *SMAD3* Knockdown Assay

Differentiated podocytes were maintained in CultureBoost-R (Cell Systems SF-4Z0-500-R) for 24 h before transfection. Based on the manufacturer’s protocol, Silencer Select siRNA for SMAD3 (Thermo, 4392420) or scramble (Thermo, 4390843) knockdown experiments were performed. Briefly, 1.5 µL Lipofectamine 3000 (Thermo, L3000001) was mixed with 25 µL Opti-MEM media (Thermo, 11058021) and incubated for 5 minutes. Next, 50 nM of siRNA or scramble was mixed with 1 µL of p3000 reagent and 25 µL Opti-MEM and incubated for 5 minutes. The two solutions were mixed carefully and incubated at room temperature for 15 minutes. 50 µL of the mixture was added to each well of podocytes. The media was swirled gently to ensure uniform siRNA or scramble distribution throughout the well and incubated at 37 °C with 5% CO_2_. After treatment, the cells were lysed for protein isolation and western blot analyses.

### Differentiation of posterior mesoderm cells

Human iPS cells were dislodged using enzyme-free dissociation buffer (Gibco, 13150-016) and centrifuged twice at 200 g for 5 min each in advanced DMEM/F12 (Gibco; 12634010) to remove residual basement membrane and mTeSR1 media. The supernatant was aspirated, and the cell pellet was resuspended in posterior mesoderm induction media consisting of DMEM/F12 with GlutaMax (GIBCO, 10565018) supplemented with 3 μM CHIR99021 (Stemgent, 04-0004), 10 μM Y27632 (TOCRIS, 1254), and 1× B27 serum-free supplement (GIBCO, 17504044). Cells were plated on laminin-511-E8-coated plates (Takara; T-304) at a seeding density of 150,000 cells per well of a 12 well plate. The cells were cultured for 2 days with daily media change and then harvested for experiments.

### Differentiation of endothelial cells

Human iPS cells were harvested with Accutase (Thermo/Life Technologies; A1110501) and reseeded on Matrigel-coated 6-well plates at a cell density of 420,000 cells/well. After 1 day incubation, lateral mesoderm was induced using a defined medium consisting of N2B27 media, which contains neurobasal media (Invitrogen, 21103049) and DMEM/F12 glutamax (Invitrogen) in 1:1 ratio with N2 (100×) (GIBCO, 21103049) and B27 without Vitamin A (GIBCO, 12587010). The media was then supplemented with 8 µM CHIR99021and 25 ng/mL hBMP4 (VWR International LLC, 10273-372) for 3 days. On Day 4, media was replaced with chemically defined endothelial induction media consisting of StemPro-34 SFM media (GIBCO, 10639011) supplemented with Glutamax (GIBCO, 35050061) in 100:1 ratio, 2 µM forskolin (Abcam Inc., ab120058), and 200 ng/mL VEGF165. Cells were fed every day, and cultured/conditioned media was collected for endothelial expansion media preparation. On Day 7, differentiated cells were dislodged with cold Accutase treatment for 5 mins and MACS-sorted (MidiMACS™ Separator) to harvest CD144^+^ and CD31^+^ cell populations. Purified cell populations were expanded in endothelium maintenance media consisting of conditioned media diluted at 1:1 ratio with StemPro-34 SFM supplemented with 2 µg/mL heparin (STEMCELL Technologies, 07980). Media was replaced every other day or until the conditioned media was depleted. For continued expansion beyond the first passage, the cells were fed with viEC maintenance media prepared with StemPro-34 supplemented with 10% HI-FBS (Invitrogen, 10082147), 2 µg/mL heparin, and 50 ng/mL VEGF165 on alternate days. For long term use, the cells were passaged every 5 days using 0.25% trypsin-EDTA (Gibco; 25200-056) and plated on freshly prepared fibronectin-coated (Thermo; 356008) T-75 flasks at a seeding density of 400,000 cells per flask.

### Quantitative real-time PCR

Cells were washed with dPBS (Gibco; 14190144) and lysed with RA1 buffer (Macherey-Nagel; 740955.250). RNA was extracted using the NucleoSpin RNA kit (Macherey-Nagel; 740955.250) following the manufacturer’s instructions. RNA (1 µg) was reverse transcribed into cDNA libraries using Superscript III RT-PCR kit (Invitrogen; 18080-093), and qPCR was performed with QuantStudio3 (Applied Biosystems) using qPCR SYBR Master Mix (Promega; A6001) to detect SYBR Green 1. Fold change was calculated using the ΔΔC_t_ method, in which samples were internally normalized to GAPDH housekeeping gene.

### Western Blot Analysis

For collection of whole cell lysates, the cells were washed twice with ice-cold dPBS (Gibco; 14190144) and lysed on ice in RIPA buffer (Millipore/Sigma; R0278-500ML) supplemented with PhosSTOP phosphatase inhibitors (Millipore/Sigma; 04906837001) and complete EDTA-free protease inhibitor cocktail (Millipore/Sigma; 4693132001). One tablet each of the PhosSTOP phosphatase inhibitor and complete EDTA-free protease inhibitor were added in 10 mL RIPA Buffer. Protein samples were separated by SDS-PAGE using Mini-PROTEAN TGX Stain-Free Precast 4-15% Gels (Bio-rad; 4568083) and transferred onto a PVDF membrane (Bio-rad; 1704157) using a Trans-blot Turbo semi-dry transfer system (Biorad; 1704150). Membranes were blocked with 5% Blotto (ChemCruz; sc-2324) in Tris-buffered saline with Tween 20 (TBST), and immunoblotting was carried out according to standard procedure. Primary antibodies were diluted in TBST supplemented with 5% Blotto and incubated for 1 h at room temperature on a platform angle rocker. Primary antibodies used for Western blot were rabbit Anti-Phospho-SMAD1 (Ser206) (Cell Signaling; mAb #5753, 1:1000), rabbit Anti-SMAD2 (D43B4) (Cell Signaling; mAb #5339, 1:1000) rabbit Anti-SMAD3 (C67H9) (Cell Signaling; mAb #9523, 1:1000), rabbit Anti-Phospho-SMAD3 (C25A9) (Cell Signaling; mAb #9520, 1:1000), rabbit Anti-SMAD4 (D3R4N) (Cell Signaling; mAb #46535, 1:1000), rabbit Anti-SMAD6 (AbClonal; pAb #9520, 1:250), rabbit Anti-SMAD7 (AbClonal; pAb #16396, 1:250), rabbit Anti-Brachyury (Abcam; ab20680, 1:1000), rabbit Anti-CDX2 antibody (Abcam; ab76541, 1:1000), goat Anti-Goosecoid (R&D Systems, AF4086, 1:500), rabbit Anti-PAX2 (Cell Signaling; mAb #9666S, 1:1000), rabbit Anti β-Catenin (D10A8) (Cell Signaling; mAb #8480, 1:1000), mouse Anti-Twist antibody (Abcam; ab175430, 1:1000), rabbit Anti-Snail (C15D3) (Cell Signaling; mAb #3879, 1:1000), rabbit Anti-alpha smooth muscle Actin antibody (Abcam; mAb #ab7817, 1:500), rabbit Anti-MLKL antibody (Abcam; mAb #ab184718, 1:500), guinea pig Anti-Nephrin antibody (ARP; #GPN02, 1:500), rabbit Anti-Podocin (Abcam, mAb #ab50339, 1:1000), mouse Anti-Synaptopodin (D-9) (SantaCruz, mAb #sc-515842, 1:1000), mouse Anti-CD2AP (B-4) (SantaCruz, mAb #sc-25273, 1:1000), mouse Anti-GLEPP1 (B-6) (SantaCruz, mAb #sc-365354, 1:1000), mouse Anti-YAP (63.7) (SantaCruz; mAb #sc-101199, 1:1000), rabbit Anti-phospho-YAP (D9W2I) (Cell Signaling; mAb #13008, 1:1000), rabbit Anti-TEAD1 (D9X2L) (Cell Signaling; mAb #12292, 1:1000), rabbit Anti-CTGF (D8Z8U) (Cell Signaling; mAb #86641, 1:1000), rabbit Anti-Naked2 (C67C4) (Cell Signaling; mAb #2073, 1:1000), mouse Anti-VE-Cadherin (BV-9) (SantaCruz, mAb #sc-52751, 1:1000); rabbit Anti-transgelin (AbClonal, pAb, #A6760, 1: 1000), rabbit Anti-ICAM1/CD54 (AbClonal, pAb, #A5597, 1:1000), rabbit Anti-vimentin (AbClonal, pAb, #A2584, 1:1000), sheep Anti-PECAM1 (Bio-Rad, mAb, #CO.3E1D4, 1:1000), and rabbit Anti-GAPDH (Millipore Sigma; #ABS16, 1:10000). Secondary antibodies --HRP-conjugated goat anti-mouse (Cell Signaling; #7076, 1:5000),HRP-conjugated goat anti-rabbit (Cell Signaling; #7074, 1:10000), HRP-conjugated goat anti-guinea pig polyclonal (ARP; #90001, 1:1000), and HRP-conjugated donkey anti-goat polyclonal (R&D Systems, #HAF109, 1:1000) -- were diluted in TBST supplemented with 5% Blotto and incubated for 1 h at room temperature on a platform angle rocker. Chemiluminescence was detected using the Super Signal West Femto kit (Thermo; 34094). Signal intensities were analyzed using a ChemiDoc imager (Bio-rad).

### Flow Cytometry

Cells were dissociated with 0.25% trypsin-EDTA (Gibco; 25200-056) and centrifuged for 5 min at 201 *g.* Supernatant was removed and washed with 1% BSA in dPBS. The cells were resuspended in resuspension buffer according to eBioscience™ Foxp3 / Transcription Factor Staining Buffer Set manufacturer protocol (Invitrogen; #00-5523-00) followed by staining with primary antibodies. The primary antibodies used included Alexa Fluor 405-conjugated human Anti-WT1 Antibody (R&D technologies; #IC57291V), PE-conjugated Anti-SNAI1 Antibody (G-7) (SantaCruz; #sc-271977 PE), Recombinant Alexa Fluor 488-conjugated Anti-MLKL antibody (Abcam; #ab207901), and Alexa Fluor 647-conjugated Anti-Pax2 antibody (Abcam; #ab218058). Cells were analyzed with a F01: BD LSRFortessa Cell Analyzer and the data was analyzed with FlowJo V10 (FlowJo, LLC) software. Gating was performed on the basis of singlets.

### Immunostaining and Microscopy

The cells were fixed with 4% paraformaldehyde (Sigma-Aldrich, #16005) in dPBS followed by permeabilization with 0.125% triton X-100 in dPBS for 5 min. Cells were blocked by incubation with a solution of 1% BSA and 0.125% triton X-100 in dPBS (blocking buffer) for 30 min at room temperature. The cells were incubated with primary antibodies in permeabilization buffer overnight at 4 °C. The cells were washed with permeabilization buffer three times and incubated with secondary antibodies conjugated to either Alexafluor-488 (Life Technologies; #A21202 1:1000) or Alexafluor-594 (Life Technologies; #A21203 1:1000) in permeabilization buffer for 1 h at room temperature. Following incubation, the cells were washed three times with permeabilization buffer and counterstained with 4′,6-diamidino-2-phenylindole (DAPI) (Invitrogen #D1306). The primary antibodies used included rabbit Anti-Brachyury (Abcam; ab20680, 1:250), rabbit Anti-CDX2 antibody (Abcam; ab76541, 1:250), goat Anti-Goosecoid (R&D Systems, AF4086, 1:250), mouse Anti-Twist antibody (Abcam; ab175430, 1:500), rabbit Anti-alpha smooth muscle Actin antibody (Abcam; mAb #ab7817, 1:500), mouse Anti-WT1 (6F-H2) (Millipore Sigma #05-753, 1:250), mouse Anti-CD2AP (B-4) (SantaCruz, mAb #sc-25273, 1:1000), mouse Anti-GLEPP1 (B-6) (SantaCruz, mAb #sc-365354, 1:250), rabbit Anti-PAX2 (EP-3251) (Abcam, mAb #ab79389, 1:250), mouse Anti-GLEPP1 (B-6) (SantaCruz, mAb #sc-365354, 1:1000), mouse Anti-YAP (63.7) (SantaCruz; mAb #sc-101199, 1:250), guinea pig Anti-Nephrin antibody (ARP; #GPN02, 1:250), rabbit Anti-Podocin (Abcam, mAb #ab50339, 1:250), mouse Anti-Synaptopodin (D-9) (SantaCruz, #sc-515842, 1:250), VE-cadherin (SantaCruz, #sc-9989, 1:200), PECAM (R&D Systems, AF-806, 1:200), rabbit Anti-transgelin (AbClonal, pAb, #A6760, 1: 250), and rabbit Anti-ICAM1/CD54 (AbClonal, pAb, #A5597, 1:250). Actin cytoskeletal structures were visualized by staining withAlexa Fluor 594 Phalloidin (Invitrogen, A12381, 1:1000). Immunostained cells were visualized with a Zeiss 880 inverted confocal Airyscan microscope and Zeiss 780 Upright confocal microscope equipped with 63x/1.4 Oil Zeiss Plan-Apochromat 44 07 62 (02) WD 0.19 mm objective. Data was analyzed using the Zen 2.3 Black software. The phase contrast images were captured using an EVOS M7000 microscope.

For live microscopy experiments, iPS cell-derived IM cells were cultured on laminin E8-coated 12-well plates. The differentiating IM cells were cultured in IM differentiation medium described above supplemented with 1X Penicillin-Streptomycin-Glutamine (GIBCO, 10378016). Differential Interference Contrast (DIC) images were acquired on a Zeiss Axio Observer Z1 Microscope equipped with an EMCCD camera, zero-drift compensation reflection-based autofocus, Pecon XL S1 incubator and control modules with heating at 37 °C and 5% humidity control, ASI Motorized stage with linear encoding, IX81 motorized stand, and **10x**/0.30 Plan-NeoFluar Ph1 (440331-9902), WD: 5.2mm. The microscope setup was controlled by MetaMorph Software 7.8 system ID 8562 (Molecular Devices). Images were automatically acquired at 1-h intervals for 167 h in a HP Z4 G4 Workstation (Intel W-2133 3.6GHz processor). Migration density was calculated using the FIJI plugins TrackMate and Chemotaxis. Migration density was calculated using the FIJI plugins TrackMate and Chemotaxis. TrackMate was used to track the cells for each cell line, and Chemotaxis plotted the TrackMate migration data as a rose plot. Briefly, a 400x400 pixel image size was used as a sample in each rose plot. A Laplacian of Gaussian (LoG) detector was used to identify the cells because it allows for the detection of objects of a specified radius. A representative cell was measured to be 30 pixels, so 30 was used as the object diameter in the LoG search. Approximately 200 spots were filtered out for each timelapse, and the spots were used to generate tracks. The Simple Linear Assignment Problem (Simple LAP) tracker was then implemented to assign the spots to separate tracks representative of cells. Cells were tracked throughout the differentiation timeline across 3 biologically independent experiments to calculate migration density. The generated tracks were represented in the form of a rose plot. Next, we manually parsed out the track number, timestamp, x-coordinate, and y-coordinate. Once the rose plots were generated, the corresponding data table was used to obtain the lengths of each sector petal (10 degrees each).”

### Single-cell data visualization

The gene expression profiles of normal and diseased human podocytes were obtained from the CZ CELLxGENE Explorer (https://cellxgene.cziscience.com). Briefly, five studies were chosen based on the available database, and the results were grouped based on disease types (AKI and CKD)^103–107^. The differential gene expression pattern was generated as a bubble plot. Genes of interest were selected based on the current work.

### Cell viability assay

Quantification of cell viability was carried out using the Cell-Counting kit 8 assay (CCK-8, Millipore-Sigma; 96992) following manufacturer’s instructions. Briefly, the CCK-8 reagent was diluted 1:10 in CultureBoostR and incubated with the cells at 37 °C and 5% CO_2_ for 2 h. The incubated mixture was then transferred to a clear, flat-bottom 96-well plate (Nunc; #269620), and the absorbance was measured at 450 nm using a FLUOstar Optima plate reader (BMG Labtech). Media blank was subtracted from the raw OD values to calculate percent viability and then normalized to signals from the WT podocytes.

### Click-iT EdU incorporation Assay

Podocyte proliferation/DNA replication was analyzed using the Click-iT™ EdU Cell Proliferation Kit with Alexa Fluor™ 594 dye (Invitrogen, C10339) following the manufacturer’s protocol. Briefly, podocytes were incubated with 10 μM EdU solution in podocyte induction medium for 24 h at 37 °C. The cells were fixed with 4% paraformaldehyde for 20 min at room temperature, washed with 3% BSA in PBS, and then permeabilized by treatment with 0.5% triton X-100 for 20 min at room temperature. The cells were incubated with the Click-iT reaction cocktail for 30 min followed by DAPI counterstaining. Cells were visualized using a EVOS M7000 microscope, and images were analyzed using Fiji (version 2.1.0/1.53c).

### ELISA

Human Activin A ELISA (Invitrogen, EHACTIVINA) was performed following the manufacturer’s instructions. Briefly, cell culture supernatant was diluted 10-fold in Assay Diluent, and 100 µL of the solution was added to the wells. The wells were covered and incubated at room temperature for 2.5 h in the dark with gentle shaking. The solution was discarded, and the wells were washed four times with 1x Wash Buffer. Next, 100 µL of prepared solution of biotin conjugate was added to the wells, and plates were incubated at room temperature for 1 h with gentle shaking. The solution was discarded, and the wells were washed four times with 1X Wash Buffer. Streptavidin-HRP solution (100 µL) was added, and plates were incubated at room temperature for 45 min with gentle shaking followed by washing with 1X Wash Buffer. Next, 100 µL TMB solution was added to the wells, and the color changed to blue. Streptavidin-HRP solution (100 µL) was added, and plates were incubated at room temperature for 30 mins in the dark with gentle shaking. Stop solution (50 µL) was added to the wells, and the color changed to yellow from blue. The absorbance was read immediately at 450 nm. Prism 9 (Version 9.5.1) software was used to interpolate the standard curve, and the background absorbance (from the media) was subtracted from the standards and the samples.

### Glomerulus-on-a-chip tissue modeling

Organ-chip microfluidic devices (10231-2; LOT 000568) were purchased from Emulate Bio. Inc. (Boston, MA, USA). The chips were activated in a plasma etcher (Emitech K-1050X) by treatment with oxygen plasma (100 W, 0.8 mbar, 30 s). The activated chips were immediately coated with 50 μg/mL laminin-511 solution (BioLamina, LN511-0502) in PBS with calcium and magnesium in their fluidic channels -- urinary (apical) and microvascular (basal) channels (separated by a porous PDMS membrane) and incubated overnight at 37 °C with 5% CO_2_. The fluidic channels were washed with advanced DMEM/F12 (Gibco; 12634010), seeded with differentiated endothelial cells (9 × 10^4^ cells in ∼15 µL) in the basal channel of the chips, and incubated by inversion to enable endothelial cell attachment on the porous membrane separating the two channels. After 5 h, the basal channel was washed with warm CultureBoost-R (Cell Systems SF-4Z0-500-R) to remove unbound cells, and the chips were incubated overnight at 37 °C with 5% CO2. The next day, differentiated IM cells (both mutant and WT, in separate chips; 1.5 × 10^5^ cells in ∼30 µL) were seeded in the apical channel and incubated for 5 h. The attached cells were then continuously perfused with their respective cell-culture media using an Ismatec IPC-N digital peristaltic pump (Cole-Parmer) at a volumetric flow rate of 246 μl h^−1^ (shear stress of 4.09e-3 dyn cm^−2^ for the top channel and 0.07 dyn cm^−2^ for the bottom channel). Cell culture media was recirculated and recirculated media was replaced every alternate day from the falcon tubes (**Extended Data** Fig. 6b). After 4 days of podocyte differentiation, the podocyte induction media was replaced with CultureBoost-R, and the resulting glomerulus chip was maintained for an additional 7 days. Spent medium was collected from the apical outlet reservoir of the glomerulus chips after completion of the podocyte differentiation for downstream analysis. For immunostaining, the chips were fixed, permeabilized, and blocked as described above. Immunostained chips were imaged with a Leica SP8 Upright Confocal Microscope, using a 25×/0.95 HCXIRAPO water dipping lens with 2.4 mm DIC objective, and Olympus FV3000-RS confocal microscope, using a 30× silicon immersion objective (0.8 MM WD, 1.05 NA). Images were sampled by the Nyquist criterion at the full resolution (i.e., NA) of the lens. The full dynamic range of the detector was maximized in image acquisition. Images were processed using CellSens software version 2.4.1. and Fiji software version 2.1.0/1.53c. Each chip was used as a biological repeat for one experiment. One-way analysis of variance (ANOVA) was performed to determine statistical significance, as indicated in the figure legends. Podocyte arborizations at the podocyte-endothelial cell interface was analyzed using the 3D viewer plugin of Fiji software (version 2.1.0/1.53c).

### Barrier function/glomerular filtration analysis

Immediately after completion of podocyte induction, CultureBoost-R media supplemented with a mixture of 10 μg/ml inulin conjugated to FITC (Sigma-Aldrich, F3272) and 100 μg/mL albumin conjugated to Alexa Fluor 555 (Thermo Fisher Scientific; A34786) was perfused through the microvascular channel. Outflow media was collected from the apical channel outlet, and fluorescence intensity was measured using a SpectraMax Fluorescent Plate reader (SpectraMax i3x, Molecular Devices). The amount of Inulin and Albumin filtered from the basal to the apical channel was analyzed using an equation for renal clearance: ([U] × UV)/[P]), where [U] is urinary concentration of albumin, UV = volume of media collected from the apical channel outlet reservoir, and [P] = dosing concentration in the basal channel.

### Statistical analyses

Statistical analyses were performed in Prism GraphPad 9.5.1. Statistical tests used two-tailed one-way ANOVA and post-hoc tests for multiple comparisons, or Student’s *t*-test for comparison between two groups. *P* < 0.05 was considered significant unless otherwise noted. Additional statistical information is provided in the figure legend.

## Acknowledgments

S.M. is a recipient of the Whitehead Scholarship in Biomedical Research, a Chair’s Research Award from the Department of Medicine at Duke University, a MEDx Pilot Grant on Biomechanics in Injury or Injury Repair, a Burroughs Wellcome Fund PDEP Career Transition Ad Hoc Award, a Duke Incubation Fund from the Duke Innovation & Entrepreneurship Initiative, a Genentech Research Award, and a George M. O’Brien Kidney Center Pilot Grant (P30 DK081943), and NIH Director’s New Innovator Award (Award Number DP2DK139544) which supported the study. R.B. is a recipient of the Lew’s Predoctoral Fellowship in the Center for Biomolecular and Tissue Engineering (CBTE) at Duke University (T32 Support NIH Grant T32GM800555). We acknowledge the National Heart, Lung, and Blood Institute Pediatrics Cardiac Genomics Consortium (PCGC) investigators (R01 HL151257 (C.E.S), 1UM1HL098166 (J.G.S) and others) for their support and expertise. T.W. is a recipient of Ruth L. Kirschstein National Research Service Award (NRSA) T32 Fellowship (2T32 HL 7208-46 A1). J.G.S is supported by Foundation Leducq 16 CVD 03 and C.E.S is supported by the Howard Hughes Medical Institute. We thank Dr. Derek W. Cain from the Duke Human Vaccine Institute (DHVI) Flow Cytometry Facility for assistance with the Flow Cytometry experiments and data analyses; Dr. James ‘Bo’ Faust from Evident for helping with glomerulus-on-a-chip microscopy; Dr. Yasheng Gao for assistance with live microscopy set up; Xingrui Mou for plasma treating the microfluidic organ chips; and Yasmin Roye for technical assistance with endothelial cell differentiation. The authors also thank all members of the Musah Lab and Dr. Marcie Pachino from the Graduate Communication Center (GCC) at Duke University for helpful comments on the manuscript.

## Author Contributions

S.M., T.W. and J.G.S. conceived the idea for this work. R.B., T.D.K. and S.M. designed the experiments and overall strategy for this study. T.W. performed CRISPR/Cas9 experiments and generated the SMAD2 mutant human iPS cell lines used in this study. R.B. performed the experiments including stem cell differentiation, qPCR analyses, western blots, microscopy, and the microfluidic organ chip experiments with support from the co-authors. T.D.K. performed protein quantification, endothelial cell differentiation, and EdU analysis. T.D.K. and R.B. performed CCK8 assay. A.M. analyzed the cell migration data generated by R.B. and provided new strategies for visualizing the data. S.L. and R.B. performed qPCR analyses and editing of the initial manuscript draft. R.B. and H.A. performed western blot analyses. R.B., T.D.K., and S.M. interpreted the results. R.B. wrote the original draft of the manuscript with input from the other authors. S.M. edited the manuscript. T.W. was supervised by J.G.S. and C.E.S.; All other authors were supervised by S.M.

---

**Extended Data Fig. 1:**
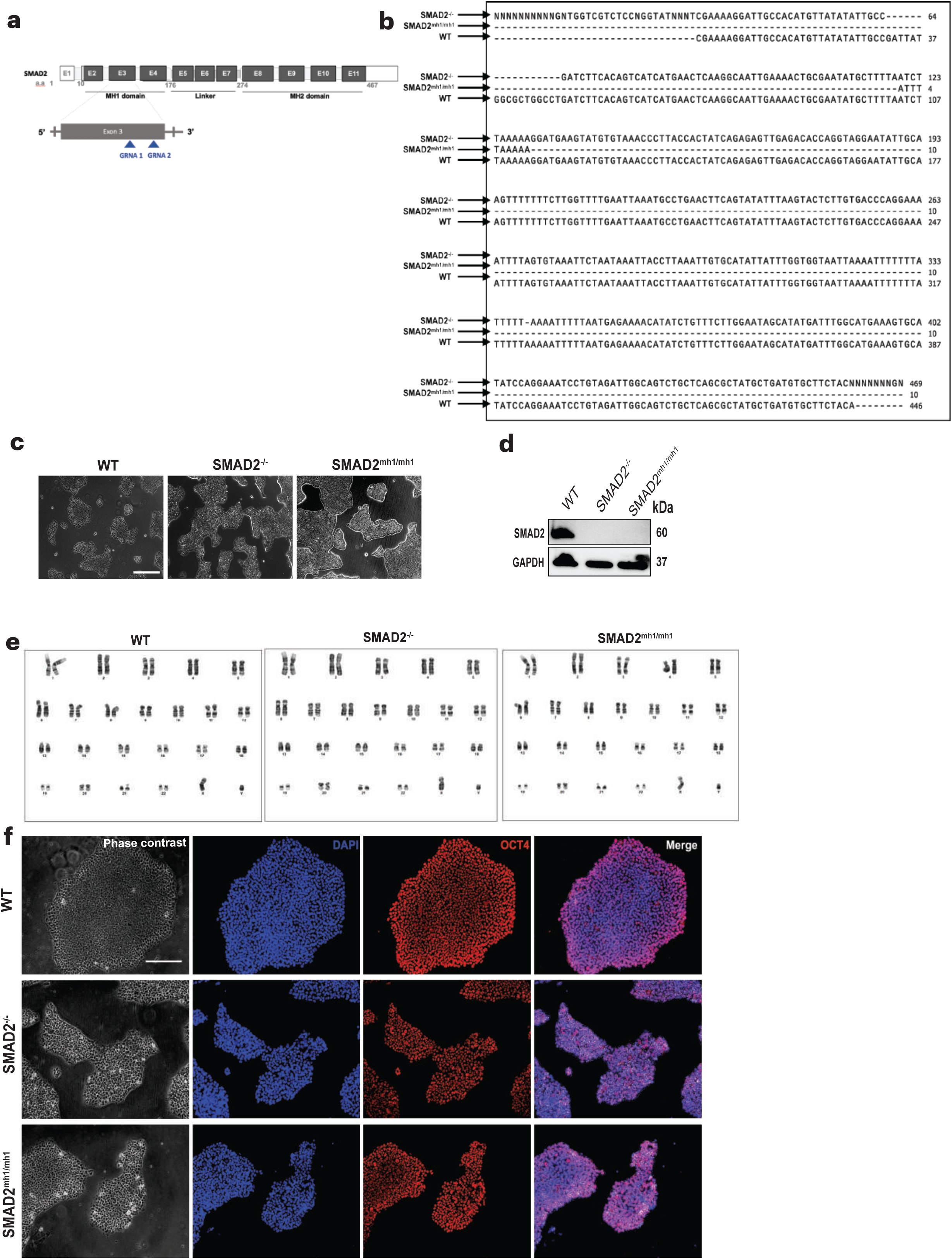
Derivation and characterization of *SMAD2* mutant cell lines. **a**, Schematic representation of *SMAD2* mutation installation in human iPS cells. **b**, Introduction of *SMAD2* variants into human iPS cells was confirmed by Sanger Sequencing. **c**, Representative phase contrast images of WT, *SMAD2*^-/-^, and *SMAD2^mh1/mh1^* iPS cells. Scale bar, 650 µm. **d**, Representative immunoblots showing loss of *SMAD2* protein in the mutant cells. **e**, Representative genetic karyotypes from the mutant cell lines after 4 passages post-gene edits. n = 20 cells from each cell line. **f**, Representative phase contrast and immunofluorescent images showing the expression of the pluripotency marker OCT4 *(red*). DAPI (blue) labels the nuclei, and the scale bar represents 150 µm.

**Extended Data Fig. 2:**
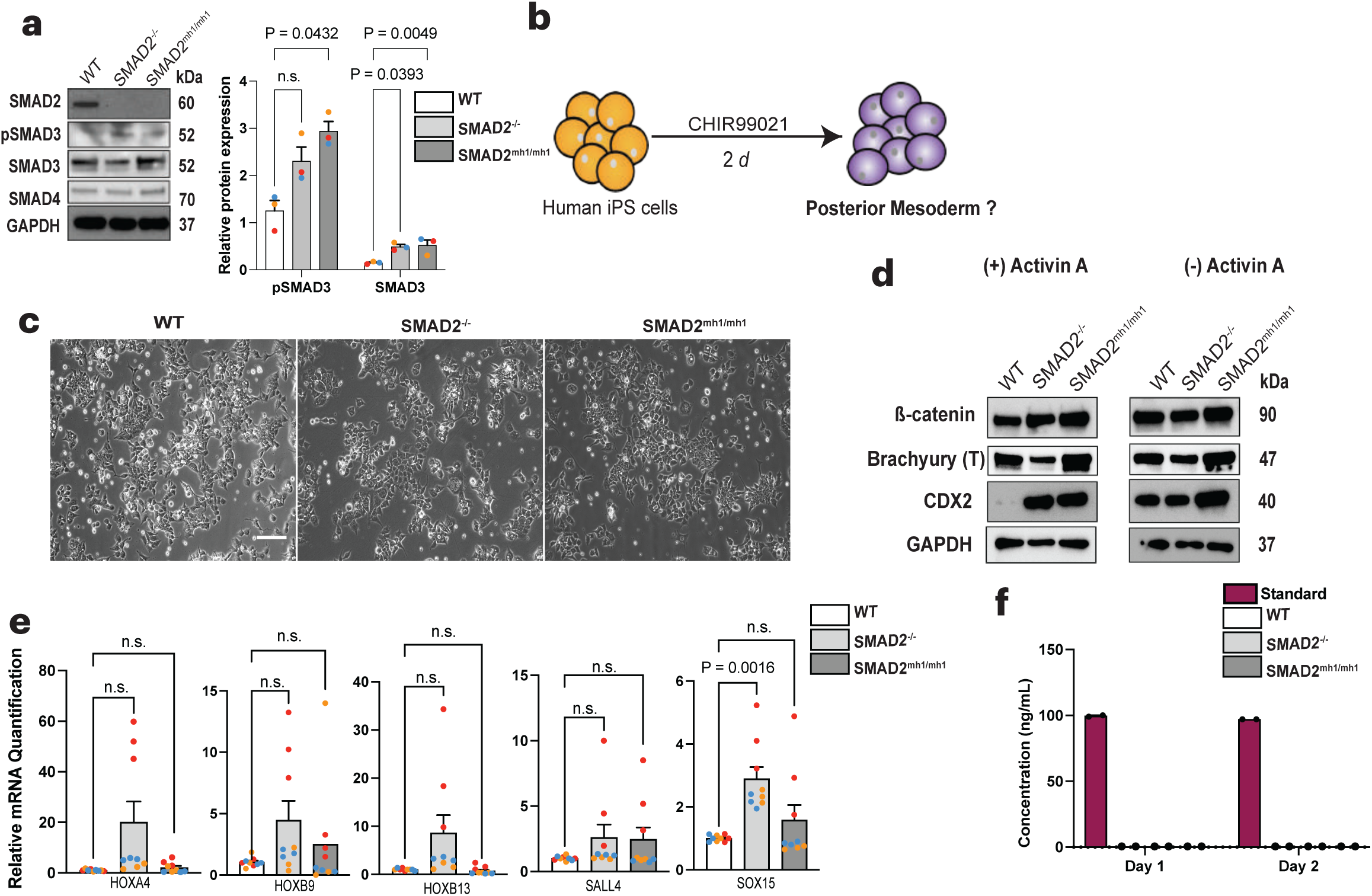
Mesoderm derivation. **a**, Representative immunoblots showing expression of *SMAD2, SMAD3*, *pSMAD3*, and *SMAD4* in the mesoderm cells. Quantifications represents mean ± SEM; n = 3 independent experiments. **b**, Schematic representation of derivation of posterior mesoderm cells with CHIR99021 pulse. **c**, Representative phase contrast images of WT, *SMAD2*^-/-^, and *SMAD2^mh1/mh1^* iPS cell-derived mesoderm generated using only CHIR99021 pulse. Images were captured after 2 days of induction. Scale bar, 275 µm. **d**, Representative immunoblots showing expression of mesoderm lineage-specific markers in the differentiated WT, *SMAD2*^-/-^, and *SMAD2^mh1/mh1^* mesoderm derived with or without exposure to Activin A. *ß-catenin* expression was slightly higher in the *SMAD2*^-/-^ and *SMAD2^mh1/mh1^* mesoderm compared to WT when mesoderm cells were differentiated with Activin A and CHIR99021. Conversely, *ß-catenin* expression was consistent in the WT, SMAD2^-/-^, and SMAD2^mh1/mh1^ mesoderm differentiated only with CHIR99021 pulse. CDX2 was expressed in WT and the mutants after treatment with CHIR99021, indicating that GSK3ß inhibition is sufficient for the generation of posterior mesoderm cells. n = 2 independent experiments. **e**, RNA expression of posterior mesoderm-specific genes in CHIR99021-treated mesoderm cells. Data are expressed relative to WT after normalizing to *GAPDH.* Data are mean values ± SEM; n = 3 independent experiments. *P*-valus were calculated by one-way ANOVA with post hoc Dunnett’s test. **f**, Quantification of cell-secreted Activin A on Days 1 and 2 during posterior mesoderm induction via CHIR99021 stimulation. The differentiating mesoderm cells did not secrete Activin A. Data are mean values ± SEM; n = 2 independent experiments.

**Extended Data Fig. 3:**
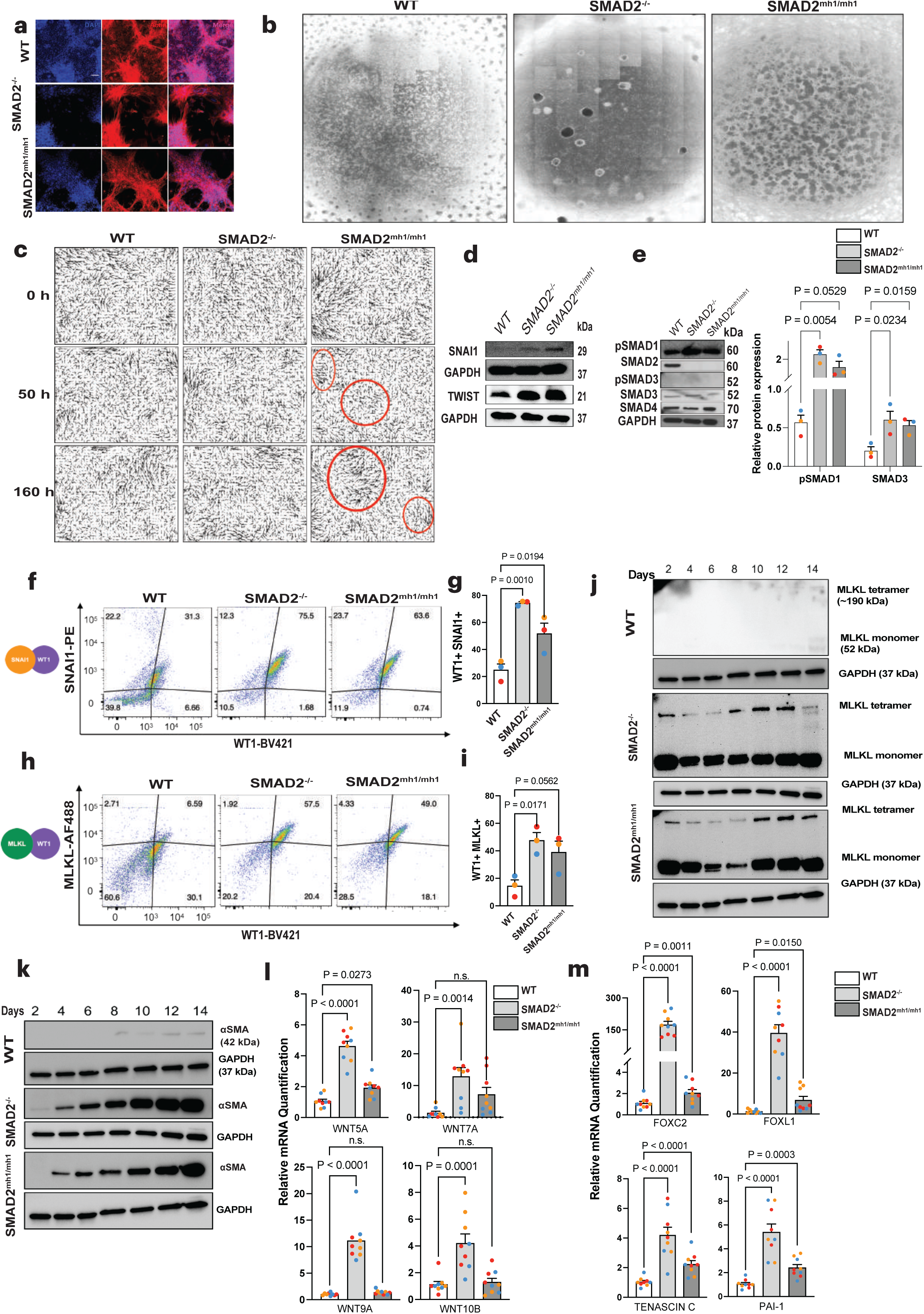
IM derivation. **a**, Representative fluorescent images of Day 14 WT, *SMAD2*^-/-^, and *SMAD2^mh1/mh1^* IM stained for Actin (with Phalloidin) and cell nuclei (DAPI). **b**, Representative whole-well scan of differentiated IM cells after 14 d of induction. WT cells grew and differentiated uniformly, *SMAD2*^-/-^ IM cells were migratory and necroptotic, and *SMAD2^mh1/mh1^*IM cells formed a honeycomb network with jammed cell clusters. **c**, Snapshots of the velocity field obtained from Particle Image Velocitometry analysis of divergence of WT and mutant IM cells. The arrows indicate local divergence to the mean direction of migration. **d**, Representative immunoblots showing expression of EMT markers *SNAI1* and *TWIST* in IM cells, which was upregulated in the mutants compared to WT. n = 2 independent experiments. **e**, Representative immunoblots showing expression of *pSMAD1, SMAD2, SMAD3*, and *SMAD4* in the IM cells. n = 3 independent experiments. **f**, Flow cytometry analysis of EMT marker *SNAI1* (PE-*SNAI1*) and BV421-*WT1*. **g**, Quantification of (PE-*SNAI1*) and (BV421-*WT1*) positive cells. Data are mean values ± SEM; n = 3 independent experiments; *p*-values were calculated by one-way ANOVA with post hoc Dunnett’s test. **h**, Flow cytometry plot showing co-expression of necroptosis marker *Mixed Lineage Kinase Domain Like Pseudokinase (MLKL)* (AF488-*MLKL*) and *WT1* (BV421-*WT1*). **i**, Quantification of (BV421-*WT1*) and (AF488-*MLKL*) positive cells. Data are mean values ± SD; n = 3 independent experiments; *p*-values were calculated by one-way ANOVA with post hoc Dunnett’s test. **j**, Representative immunoblots showing expression of *MLKL* oligomers in differentiating IM cells. A time-dependent increase in *MLKL* tetramerization in the mutant IM cells can be observed. n = 2 independent experiments. **k**, Representative immunoblots showing expression of *αSMA* in differentiating IM cells every other day. No *αSMA* expression was detected in the WT IM cells, but there was a monotonic increase in *αSMA* expression in mutant IM cells. n = 2 independent experiments. **l**, RNA expression analyses for canonical and noncanonical WNT target genes in IM cells. Data are expressed relative to WT after normalizing to *GAPDH*. Data are mean values ± SEM; n = 3 independent experiments. *P-* values were calculated by one-way ANOVA with post hoc Dunnett’s test. **m**, RNA expression of mesenchymal genes in IM cells. Data are expressed relative to WT after normalizing to *GAPDH.* Data are mean values ± SEM; n = 3 independent experiments. *P*-values were calculated by one-way paired t-test two-tailed.

**Extended Data Fig. 4:**
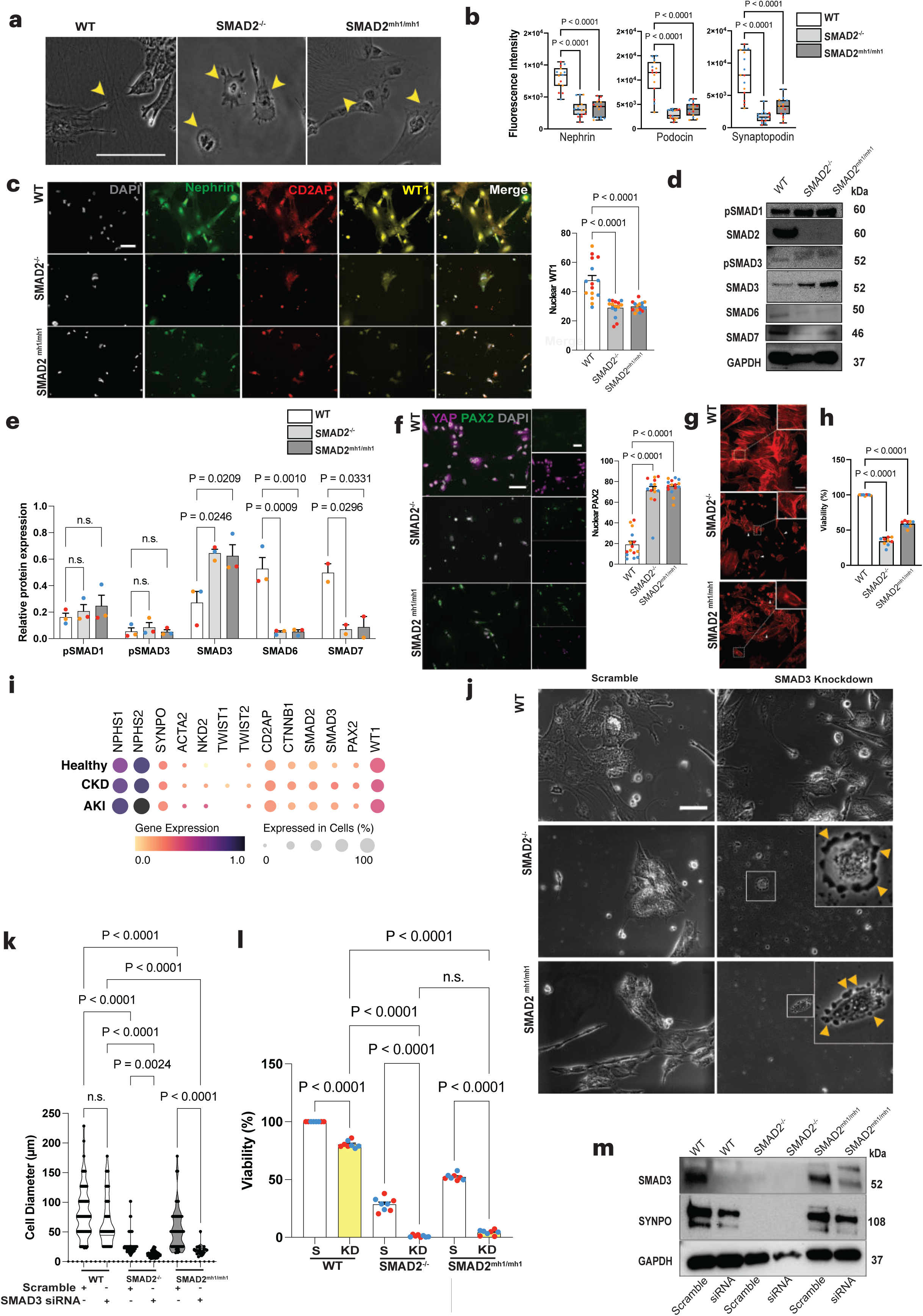
Podocyte derivation. **a**, Phase contrast images showing podocyte foot processes (WT) and effacement-like phenotype (mutants). Scale, 150 µm. **b**, Fluorescence intensity quantification of podocyte lineage specific markers nephrin, podocin, and synaptopodin. Data are mean values ± SEM; n = 12 fields of view from 2 independent experiments. Expression of nephrin, podocin, and synaptopodin were significantly reduced in the *SMAD2*^-/-^ and *SMAD2^mh1/mh1^* podocytes with respect to the WT. *p*-values were calculated by one-way ANOVA with post hoc Dunnett’s test. **c**, Representative immunofluorescent images showing the expression of *Nephrin* (green), *CD2AP* (red), and *WT1* (yellow) in differentiated podocytes. DAPI, shown in grey, labels the nuclei, and the scale bar represent 100 µm for all images. The nuclear intensity of WT1 was significantly downregulated in the mutant podocytes. Data are mean values ± SEM; n = 5 fields of view from 3 independent experiments. *P*-value was calculated using one-way ANOVA analysis with post hoc Dunnett’s test. **d**, Representative immunoblots showing expression of *pSMAD1, SMAD2, pSMAD3, SMAD3, SMAD6,* and *SMAD7* in podocytes across 3 biologically independent experiments. **e**, Relative protein expression of *SMAD* family proteins across 3 biologically independent experiments. *P*-value was calculated using one-way ANOVA analysis with post hoc Dunnett’s test. **f**, Representative immunofluorescent images showing the expression of *YAP* (magenta) and *PAX2* (green). DAPI, shown in grey, labels the nuclei, and the scale bar represent 100 µm for all images. The nuclear intensity of PAX2 was significantly increased in the mutant podocytes. Data are mean values ± SEM; n = 5 fields of view from 2 independent experiments. *P*-value was calculated using one-way ANOVA analysis with post hoc Dunnett’s test. **g**, Mutant podocytes demonstrated altered cytoskeletal rearrangement characterized by the presence of distinct local actin microdomains. Top right: zoomed-in regions, marked by dashed white boxes, indicate actin dissolution and subsequent podocyte effacement. Scale bar, 100 µm. **h**, CCK-8 cell viability assay on differentiated podocytes. Cell viability was significantly lower in the mutants compared to WT. Data are mean values ± SEM. n = 3 independent experiments. *P*-value was calculated using one-way ANOVA analysis with post hoc Dunnett’s test. **i**, Bubble plot comparing the expression of markers in healthy and diseased human kidney podocytes. Data was obtained by reanalyzing previously published literature on human kidney samples^102–106^. **j**, Representative phase contrast images of *SMAD3* knockdown podocytes. The knockdown mutant podocytes underwent foot process effacement (indicated with orange arrows) and cell death. Scale bar, 100 µm. **k**, Cell size was significantly reduced after SMAD3 knockdown. Data are mean values ± SEM. n = 3 independent experiments. *P*-value was calculated using one-way ANOVA analysis with post hoc Sidak’s test.**l**, CCK-8 cell viability assay on differentiated podocytes treated with scrambled or *SMAD3* siRNA. Cell viability was significantly reduced in the mutants compared to WT scrambled or knockdown podocytes. Data are mean values ± SEM. n = 3 independent experiments. *P*-value was calculated using one-way ANOVA analysis with post hoc Dunnett’s test. **m**, Representative immunoblots showing expression of *SMAD3* and *Synaptopodin* after *SMAD3* knockdown in podocytes. n = 2 biologically independent experiments.

**Extended Data Fig. 5:**
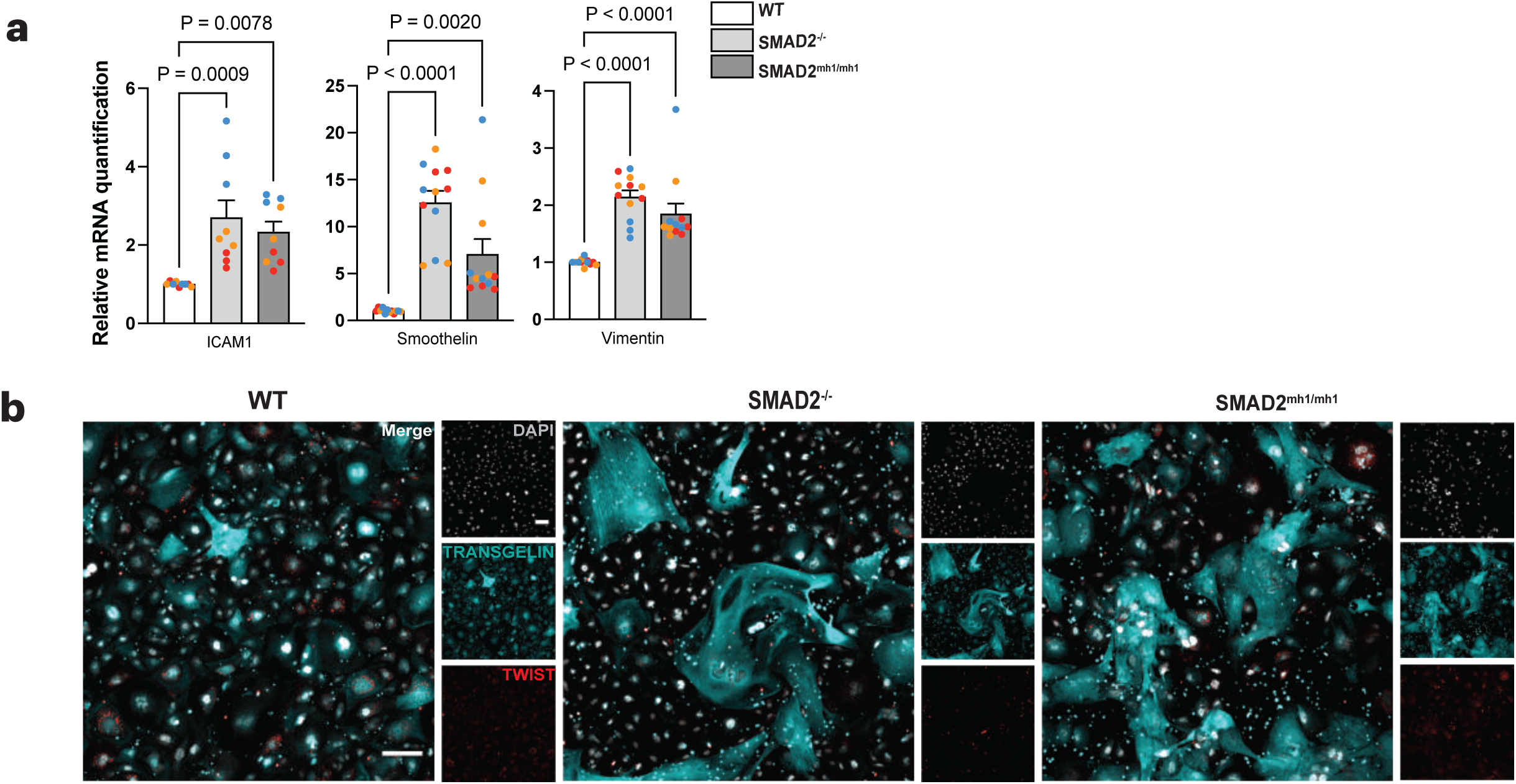
Endothelial cell derivation. **a**, RNA expression of mesenchymal markers ICAM1, Smoothelin, and Vimentin in WT and mutant endothelial cells. Data are expressed relative to WT after normalizing to *GAPDH*. Data are mean values ± SEM; n = 3 independent experiments. *P*-values were calculated by one-way ANOVA with post hoc Dunnett’s test. **b**, Representative immunofluorescent images showing the expression of mesenchymal markers *Transgelin* (cyan) and *TWIST* (red) in endothelial cells after 2 weeks of maintenance culture. Single-channel images are shown next to the merged images for the entire field of view. DAPI, shown in grey, labels the nuclei, and the scale bar represent 100 µm for all images.

**Extended Data Fig. 6:**
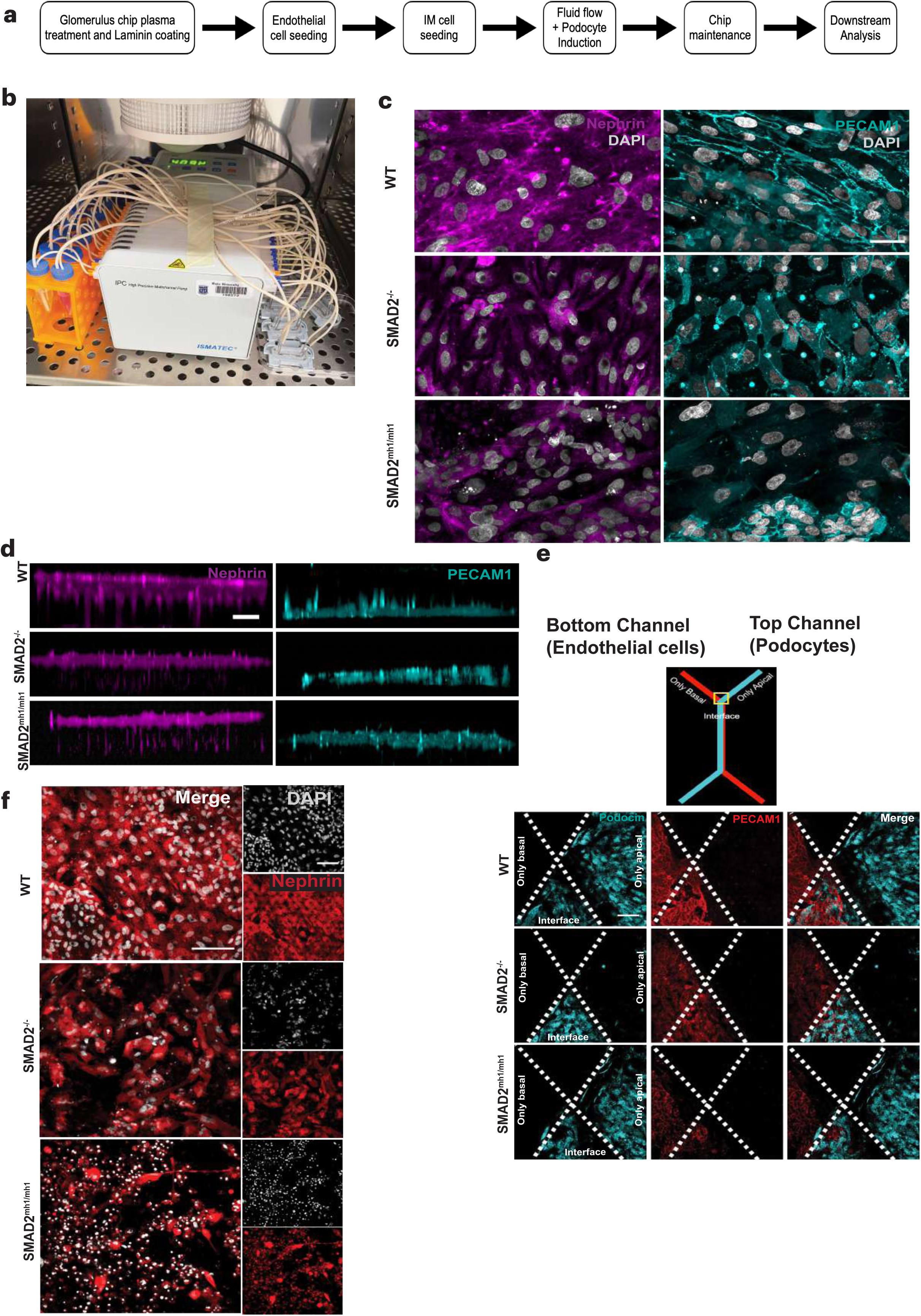
Recapitulating the glomerular capillary wall using a glomerulus-on-a-chip device. **a**, Schematic representation of the development of glomerulus-on-a-chip cultures. **b**, Example photograph of the experimental setup for glomerulus-on-a-chip cultures connected to two reservoirs for the cell culture media for the urinary (podocyte induction media) and capillary (CultureBoost-R) channels. These microfluidic glomerulus-on-a-chips were connected to media reservoirs with low retention inlet and outlet tubing via a peristaltic pump. Media was recirculated and replenished every alternate day. **c**, Representative immunofluorescence micrograph of WT and mutant glomerulus-on-a-chip showing *Nephrin (magenta)*-and *PECAM1 (cyan)*-positive podocytes and endothelial cells, respectively. **d**, Representative single channel images of *Nephrin^+^*podocytes (magenta, urinary channel) and *PECAM1^+^* endothelial cells (cyan, capillary channel) along the z axis. Scale bar, 30 µm. **e**, Representative images of the ‘Y’ junction of the chips to visualize the incoming apical and basal channels before they overlay at the interface. Mutant podocytes mostly died during differentiation when not interfaced with endothelial cells, as observed by the loss of podocin in the ‘only apical’ channel of the mutant chips. **f**, Representative immunofluorescence images of WT, *SMAD2*^-/-^, and *SMAD2^mh1^*^/*mh1*^ podocytes on the glomerulus-on-a-chip platform cultured in absence of the endothelial cells. DAPI, shown in grey, labels the nuclei. Scale bar, 100 µm.

## References

1. Loh, K. M. et al. Mapping the Pairwise Choices Leading from Pluripotency to Human Bone, Heart, and Other Mesoderm Cell Types. Cell 166, 451–467 (2016).

2. Musah, S., Bhattacharya, R. & Himmelfarb, J. Kidney Disease Modeling with Organoids and Organs-on-Chips. Annu. Rev. Biomed. Eng. 26, annurev-bioeng-072623-044010 (2024).

3. Micha, D., et al. *SMAD2* Mutations Are Associated with Arterial Aneurysms and Dissections: HUMAN MUTATION. Human Mutation 36, 1145–1149 (2015).

4. Granadillo, J. L. et al. Variable cardiovascular phenotypes associated with *SMAD2* pathogenic variants. Human Mutation 39, 1875–1884 (2018).

5. Loeys, B. L. et al. Aneurysm Syndromes Caused by Mutations in the TGF-β Receptor. n engl j med 11 (2006).

6. Cannaerts, E. et al. Novel pathogenic *SMAD2* variants in five families with arterial aneurysm and dissection: further delineation of the phenotype. J Med Genet 56, 220–227 (2019).

7. Bikbov, B. et al. Global, regional, and national burden of chronic kidney disease, 1990–2017: a systematic analysis for the Global Burden of Disease Study 2017. The Lancet 395, 709–733 (2020).

8. Morton, S. U., Quiat, D., Seidman, J. G. & Seidman, C. E. Genomic frontiers in congenital heart disease. Nat Rev Cardiol 19, 26–42 (2022).

9. Dovjak, G. O. et al. Abnormal Extracardiac Development in Fetuses With Congenital Heart Disease. Journal of the American College of Cardiology 78, 2312–2322 (2021).

10. Gabriel, G. C., Pazour, G. J. & Lo, C. W. Congenital Heart Defects and Ciliopathies Associated With Renal Phenotypes. Front. Pediatr. 6, 175 (2018).

11. Rajpal, S. et al. Association of Albuminuria With Major Adverse Outcomes in Adults With Congenital Heart Disease: Results From the Boston Adult Congenital Heart Biobank. JAMA Cardiol 3, 308 (2018).

12. Zaidi, S. et al. De novo mutations in histone-modifying genes in congenital heart disease. Nature 498, 220–223 (2013).

13. Homsy, J. et al. De novo mutations in congenital heart disease with neurodevelopmental and other congenital anomalies. Science 350, 1262–1266 (2015).

14. Scholes, G. B., Zannino, D., Kausman, J. Y. & Cheung, M. M. H. Altered in utero kidney development in newborns with congenital heart disease. Pediatr Res 85, 644–649 (2019).

15. Gupte, P. A., Vaideeswar, P. & Kandalkar, B. M. Cyanotic Nephropathy-A Morphometric Analysis: Cyanotic Nephropathy. Congenit Heart Dis 9, 280–285 (2014).

16. Dimopoulos, K. et al. Prevalence, Predictors, and Prognostic Value of Renal Dysfunction in Adults With Congenital Heart Disease. Circulation 117, 2320–2328 (2008).

17. Greenberg, J. H. et al. Kidney Outcomes 5 Years After Pediatric Cardiac Surgery: The TRIBE-AKI Study. JAMA Pediatr 170, 1071 (2016).

18. Nugent, J. et al. Assessment of Acute Kidney Injury and Longitudinal Kidney Function After Hospital Discharge Among Patients With and Without COVID-19. JAMA Netw Open 4, e211095 (2021).

19. Li, A. H. et al. Genetic architecture of laterality defects revealed by whole exome sequencing. Eur J Hum Genet 27, 563–573 (2019).

20. Hobbs, C. A. et al. Genetic Epidemiology and Nonsyndromic Structural Birth Defects: From Candidate Genes to Epigenetics. JAMA Pediatr 168, 371 (2014).

21. Rees, L., Schaefer, F., Schmitt, C. P., Shroff, R. & Warady, B. A. Chronic dialysis in children and adolescents: challenges and outcomes. The Lancet Child & Adolescent Health 1, 68–77 (2017).

22. Woolf, A. S. A molecular and genetic view of human renal and urinary tract malformations. Kidney International 58, 500–512 (2000).

23. Strong, A. et al. Expanding the phenotypic spectrum of Mendelian connective tissue disorders to include prominent kidney phenotypes. American J of Med Genetics Pt A 185, 3762–3769 (2021).

24. Billett, J., Majeed, A., Gatzoulis, M. & Cowie, M. Trends in hospital admissions, in-hospital case fatality and population mortality from congenital heart disease in England, 1994 to 2004. Heart 94, 342–348 (2008).

25. Keller, G., Zimmer, G., Mall, G., Ritz, E. & Amann, K. Nephron Number in Patients with Primary Hypertension. N Engl J Med 348, 101–108 (2003).

26. Boo, H. J. et al. The presence of simple renal cysts is associated with an increased risk of albuminuria in young adults. Korean J Intern Med 37, 425–433 (2022).

27. Sevillano, A. M. et al. Multiple kidney cysts in thin basement membrane disease with proteinuria and kidney function impairment. Clinical Kidney Journal 7, 251–256 (2014).

28. Chen, J., Ma, X., Xu, D., Cao, W. & Kong, X. Association between simple renal cyst and kidney damage in a Chinese cohort study. Renal Failure 41, 600–606 (2019).

29. Jin, S. C. et al. Contribution of rare inherited and de novo variants in 2,871 congenital heart disease probands. Nat Genet 49, 1593–1601 (2017).

30. Xiao, F. et al. Functional dissection of human cardiac enhancers and noncoding de novo variants in congenital heart disease. Nat Genet 56, 420–430 (2024).

31. Passwell, J. et al. Abnormal renal functions in cyanotic congential heart disease. Archives of Disease in Childhood 51, 803–805 (1976).

32. Burlet, A., Drukker, A. & Guignard, J.-P. Renal Function in Cyanotic Congenital Heart Disease. Nephron 81, 296–300 (1999).

33. Esch, J. J., Salvin, J. M., Thiagarajan, R. R., Del Nido, P. J. & Rajagopal, S. K. Acute kidney injury after Fontan completion: Risk factors and outcomes. The Journal of Thoracic and Cardiovascular Surgery 150, 190–197 (2015).

34. Gillesén, M. et al. Chronic kidney disease in patients with congenital heart disease: a nationwide, register-based cohort study. European Heart Journal Open 2, oeac055 (2022).

35. Anne, P., Du, W., Mattoo, T. K. & Zilberman, M. V. Nephropathy in patients after Fontan palliation. International Journal of Cardiology 132, 244–247 (2009).

36. Sharma, S. et al. Assessment of Kidney Function in Survivors Following Fontan Palliation: Kidney Function Following Fontan Palliation. Congenital Heart Disease 11, 630–636 (2016).

37. Opotowsky, A. R. et al. Estimated glomerular filtration rate and urine biomarkers in patients with single-ventricle Fontan circulation. Heart 103, 434–442 (2017).

38. Broda, C. R. et al. Renal dysfunction is associated with higher central venous pressures in patients with Fontan circulation. Congenital Heart Disease 13, 602–607 (2018).

39. Wilson, T. G. et al. Hepatic and renal end-organ damage in the Fontan circulation: A report from the Australian and New Zealand Fontan Registry. International Journal of Cardiology 273, 100–107 (2018).

40. Özsu, E., Yeşiltepe Mutlu, G., Yüksel, A. B. & Hatun, Ş. Features of Two Cases with 18q Deletion Syndrome. Jcrpe 6, 51–54 (2014).

41. Hochberg, E. P. et al. Case 11-2017 — A 61-Year-Old Woman with Leg Swelling, Back Pain, and Hydronephrosis. N Engl J Med 376, 1461–1471 (2017).

42. Turleau, C., Chavin-Colin, F., Narbouton, R., Asensi, D. & Grouchy, J. D. Trisomy 18q-. Trisomy mapping of chromosome 18 revisited. Clinical Genetics 18, 20–26 (1980).

43. Nomura, M. & Li, E. Smad2 role in mesoderm formation, left–right patterning and craniofacial development. Nature 393, 786–790 (1998).

44. Weinstein, M. et al. Failure of egg cylinder elongation and mesoderm induction in mouse embryos lacking the tumor suppressor *smad2*. Proc. Natl. Acad. Sci. U.S.A. 95, 9378–9383 (1998).

45. Faial, T. et al. Brachyury and SMAD signalling collaboratively orchestrate distinct mesoderm and endoderm gene regulatory networks in differentiating human embryonic stem cells. Development 142, 2121–2135 (2015).

46. Mendjan, S. et al. NANOG and CDX2 Pattern Distinct Subtypes of Human Mesoderm during Exit from Pluripotency. Cell Stem Cell 15, 310–325 (2014).

47. Ten Berge, D. et al. Wnt Signaling Mediates Self-Organization and Axis Formation in Embryoid Bodies. Cell Stem Cell 3, 508–518 (2008).

48. Henderson, N. C., Rieder, F. & Wynn, T. A. Fibrosis: from mechanisms to medicines. Nature 587, 555–566 (2020).

49. Musah, S. et al. Mature induced-pluripotent-stem-cell-derived human podocytes reconstitute kidney glomerular-capillary-wall function on a chip. Nat Biomed Eng 1, 0069 (2017).

50. Bhattacharya, R., Bonner, M. G. & Musah, S. Harnessing developmental plasticity to pattern kidney organoids. Cell Stem Cell 28, 587–589 (2021).

51. Lam, A. Q. et al. Rapid and Efficient Differentiation of Human Pluripotent Stem Cells into Intermediate Mesoderm That Forms Tubules Expressing Kidney Proximal Tubular Markers. JASN 25, 1211–1225 (2014).

52. Meng, X. M. et al. Smad2 Protects against TGF-β/Smad3-Mediated Renal Fibrosis. JASN 21, 1477–1487 (2010).

53. Samson, A. L. et al. MLKL trafficking and accumulation at the plasma membrane control the kinetics and threshold for necroptosis. Nat Commun 11, 3151 (2020).

54. Edeling, M., Ragi, G., Huang, S., Pavenstädt, H. & Susztak, K. Developmental signalling pathways in renal fibrosis: the roles of Notch, Wnt and Hedgehog. Nat Rev Nephrol 12, 426– 439 (2016).

55. Zeisberg, M. et al. BMP-7 counteracts TGF-β1–induced epithelial-to-mesenchymal transition and reverses chronic renal injury. Nat Med 9, 964–968 (2003).

56. Schnell, J., Achieng, M. & Lindström, N. O. Principles of human and mouse nephron development. Nat Rev Nephrol 18, 628–642 (2022).

57. Perico, L., Conti, S., Benigni, A. & Remuzzi, G. Podocyte–actin dynamics in health and disease. Nat Rev Nephrol 12, 692–710 (2016).

58. Wharram, B. L. et al. Altered podocyte structure in GLEPP1 (Ptpro)-deficient mice associated with hypertension and low glomerular filtration rate. J. Clin. Invest. 106, 1281–1290 (2000).

59. Kim, Y. H. et al. GLEPP1 Receptor Tyrosine Phosphatase (Ptpro) in Rat PAN Nephrosis. Nephron 90, 471–476 (2002).

60. Tossidou, I. et al. Tyrosine Phosphorylation of CD2AP Affects Stability of the Slit Diaphragm Complex. JASN 30, 1220–1237 (2019).

61. Guo, J.-K. WT1 is a key regulator of podocyte function: reduced expression levels cause crescentic glomerulonephritis and mesangial sclerosis. Human Molecular Genetics 11, 651– 659 (2002).

62. Kavsak, P. et al. Smad7 Binds to Smurf2 to Form an E3 Ubiquitin Ligase that Targets the TGF␤ Receptor for Degradation. Molecular Cell.

63. Fukasawa, H. et al. Down-regulation of Smad7 expression by ubiquitin-dependent degradation contributes to renal fibrosis in obstructive nephropathy in mice. Proc. Natl. Acad. Sci. U.S.A. 101, 8687–8692 (2004).

64. Chung, A. C. K. et al. Disruption of the Smad7 gene promotes renal fibrosis and inflammation in unilateral ureteral obstruction (UUO) in mice. Nephrology Dialysis Transplantation 24, 1443–1454 (2009).

65. Wang, W., Koka, V. & Lan, H. Y. Transforming growth factor-β and Smad signalling in kidney diseases: Review Article. Nephrology 10, 48–56 (2005).

66. Li, Z. et al. 3D Culture Supports Long-Term Expansion of Mouse and Human Nephrogenic Progenitors. Cell Stem Cell 19, 516–529 (2016).

67. Bonse, J. et al. Nuclear YAP localization as a key regulator of podocyte function. Cell Death Dis 9, 850 (2018).

68. Burt, M. A., Kalejaiye, T. D., Bhattacharya, R., Dimitrakakis, N. & Musah, S. Adriamycin-Induced Podocyte Injury Disrupts the YAP-TEAD1 Axis and Downregulates Cyr61 and CTGF Expression. ACS Chem. Biol. 17, 3341–3351 (2022).

69. Haley, K. E. et al. YAP Translocation Precedes Cytoskeletal Rearrangement in Podocyte Stress Response: A Podometric Investigation of Diabetic Nephropathy. Front. Physiol. 12, 625762 (2021).

70. Schwartzman, M. et al. Podocyte-Specific Deletion of Yes-Associated Protein Causes FSGS and Progressive Renal Failure. Journal of the American Society of Nephrology 27, 216– 226 (2016).

71. Holbourn, K. P., Acharya, K. R. & Perbal, B. The CCN family of proteins: structure–function relationships. Trends in Biochemical Sciences 33, 461–473 (2008).

72. Musah, S., Bhattacharya, R. & Himmelfarb, J. Kidney Disease Modeling with Organoids and Organs-on-Chips. (2024).

73. Musah, S. et al. Glycosaminoglycan-Binding Hydrogels Enable Mechanical Control of Human Pluripotent Stem Cell Self-Renewal. ACS Nano 6, 10168–10177 (2012).

74. Furukawa, K. T., Yamashita, K., Sakurai, N. & Ohno, S. The Epithelial Circumferential Actin Belt Regulates YAP/TAZ through Nucleocytoplasmic Shuttling of Merlin. Cell Reports 20, 1435–1447 (2017).

75. Aragona, M. et al. A Mechanical Checkpoint Controls Multicellular Growth through YAP/TAZ Regulation by Actin-Processing Factors. Cell 154, 1047–1059 (2013).

76. Campbell, K. N. et al. Yes-associated Protein (YAP) Promotes Cell Survival by Inhibiting Proapoptotic Dendrin Signaling. Journal of Biological Chemistry 288, 17057–17062 (2013).

77. Meliambro, K. et al. The Hippo pathway regulator KIBRA promotes podocyte injury by inhibiting YAP signaling and disrupting actin cytoskeletal dynamics. Journal of Biological Chemistry 292, 21137–21148 (2017).

78. Beck, B. et al. Different Levels of Twist1 Regulate Skin Tumor Initiation, Stemness, and Progression. Cell Stem Cell 16, 67–79 (2015).

79. Kuppe, C. et al. Decoding myofibroblast origins in human kidney fibrosis. Nature 589, 281– 286 (2021).

80. Lasagni, L., Lazzeri, E. & Shankland, S. J. Podocyte Mitosis – A Catastrophe. 13, 13–23 (2013).

81. Atchison, L. et al. iPSC-Derived Endothelial Cells Affect Vascular Function in a Tissue-Engineered Blood Vessel Model of Hutchinson-Gilford Progeria Syndrome. Stem Cell Reports 14, 325–337 (2020).

82. Roye, Y. et al. A Personalized Glomerulus Chip Engineered from Stem Cell-Derived Epithelium and Vascular Endothelium. Micromachines 12, 967 (2021).

83. Okafor, A. E., Bhattacharya, R. & Musah, S. Models of kidney glomerulus derived from human-induced pluripotent stem cells. in iPSCs in Tissue Engineering 329–370 (Elsevier, 2021). doi:10.1016/B978-0-12-823809-7.00013-X.

84. Robert, B., Zhao, X. & Abrahamson, D. R. Coexpression of neuropilin-1, Flk1, and VEGF 164 in developing and mature mouse kidney glomeruli. American Journal of Physiology-Renal Physiology 279, F275–F282 (2000).

85. Eremina, V. et al. Glomerular-specific alterations of VEGF-A expression lead to distinct congenital and acquired renal diseases. J. Clin. Invest. 111, 707–716 (2003).

86. Kalejaiye, T. D., Holmes, J. A., Bhattacharya, R. & Musah, S. Reconstitution of the kidney glomerular capillary wall. in Regenerative Nephrology 331–351 (Elsevier, 2022). doi:10.1016/B978-0-12-823318-4.00007-X.

87. Dittrich, S., et al. Renal impairment in patients with long-standing cyanotic congenital heart disease. (1998).

88. Krull, F. et al. Renal Involvement in Patients with Congenital Cyanotic Heart Disease. Acta Paediatrica 80, 1214–1219 (1991).

89. Zheng, J., Yao, Y., Han, L. & Xiao, Y. Renal function and injury in infants and young children with congenital heart disease. Pediatr Nephrol 28, 99–104 (2013).

90. van der Linde, D. et al. Birth Prevalence of Congenital Heart Disease Worldwide. Journal of the American College of Cardiology 58, 2241–2247 (2011).

91. Henry, G. L. & Melton, D. A. *Mixer* , a Homeobox Gene Required for Endoderm Development. Science 281, 91–96 (1998).

92. Nishita, M. et al. Interaction between Wnt and TGF-␤ signalling pathways during formation of Spemann’s organizer. 403, (2000).

93. Hofbauer, P. et al. Cardioids reveal self-organizing principles of human cardiogenesis. Cell 184, 3299–3317.e22 (2021).

94. Wagner, K.-D. et al. An Inducible Mouse Model for PAX2-Dependent Glomerular Disease: Insights into a Complex Pathogenesis. Current Biology 16, 793–800 (2006).

95. Dressler, G. R. et al. Deregulation of Pax-2 expression in transgenic mice generates severe kidney abnormalities. Nature 362, 65–67 (1993).

96. Murer, L. et al. Expression of Nuclear Transcription Factor PAX2 in Renal Biopsies of Juvenile Nephronophthisis. Nephron 91, 588–593 (2002).

97. Fang, N.-W. et al. Incidence and risk factors for chronic kidney disease in patients with congenital heart disease. Pediatr Nephrol 36, 3749–3756 (2021).

98. Le Jemtel, T. H. et al. Direct Evidence of Podocyte Damage in Cardiorenal Syndrome Type 2: Preliminary Evidence. Cardiorenal Med 5, 125–134 (2015).

99. Heyer, J. et al. Postgastrulation *Smad2-* deficient embryos show defects in embryo turning and anterior morphogenesis. Proc. Natl. Acad. Sci. U.S.A. 96, 12595–12600 (1999).

100. Duan, W.-J., Yu, X., Huang, X.-R., Yu, J. & Lan, H. Y. Opposing Roles for Smad2 and Smad3 in Peritoneal Fibrosis in Vivo and in Vitro. The American Journal of Pathology 184, 2275–2284 (2014).

101. Sharma, A. et al. CRISPR/Cas9-Mediated Fluorescent Tagging of Endogenous Proteins in Human Pluripotent Stem Cells. Current Protocols in Human Genetics 96, (2018).

102. Musah, S., Dimitrakakis, N., Camacho, D. M., Church, G. M. & Ingber, D. E. Directed differentiation of human induced pluripotent stem cells into mature kidney podocytes and establishment of a Glomerulus Chip. Nat Protoc 13, 1662–1685 (2018).

103. Li, R. et al. Mapping single-cell transcriptomes in the intra-tumoral and associated territories of kidney cancer. Cancer Cell 40, 1583–1599.e10 (2022).

104. Wilson, P. C. et al. Multimodal single cell sequencing implicates chromatin accessibility and genetic background in diabetic kidney disease progression. Nat Commun 13, 5253 (2022).

105. Stewart, B. J. et al. Spatiotemporal immune zonation of the human kidney. Science 365, 1461–1466 (2019).

106. Lake, B. B. et al. An atlas of healthy and injured cell states and niches in the human kidney. Nature 619, 585–594 (2023).

107. Muto, Y. et al. Single cell transcriptional and chromatin accessibility profiling redefine cellular heterogeneity in the adult human kidney. Nat Commun 12, 2190 (2021).

